# Neuronal signatures of successful one-shot memory in mid-level visual cortex

**DOI:** 10.1101/2025.09.22.677855

**Authors:** Grace F. DiRisio, Cheng Xue, Marlene R. Cohen

## Abstract

High-capacity, one-shot visual recognition memory challenges theories of learning and neural coding because it requires rapid, robust, and durable representations. Most studies have focused on the hippocampus and other higher areas. However, behavioral evidence demonstrating links between image properties and memorability and revealing image specificity of visual memory suggests an important role for mid-level visual cortex. We tested the hypothesis that area V4 contains signals that could support recognition memory. Our task increased difficulty, allowing comparisons of neuronal population responses on correct and error trials. We observed signatures of several proposed memory mechanisms including magnitude coding, repetition suppression, sparse coding, and population response consistency, but only sparse coding and population response consistency predicted behavior. Familiar images also evoked faster dynamics, consistent with pattern completion. These findings demonstrate that the building blocks of fast, high-capacity memory are present in mid-level sensory cortex, highlighting its role in distributed memory networks.

## Introduction

Humans and other primates can remember thousands of images after a single viewing, accurately reporting whether an image is novel or familiar^1,2^. This form of one-shot, high-capacity memory challenges conventional theories of learning, memory, and neural coding because it combines speed, generalization, and durability. A central question is how neuronal populations can support such a capacity.

Studying the neuronal substrates of visual recognition memory is challenging. Memories form rapidly, often after a single exposure, complicating the separation perception, encoding, and retrieval^1,3–6^. Natural images inhabit a high-dimensional feature space, making neuronal responses difficult to quantify and interpret. Trial-to-trial response variability, which has been implicated in attention, working memory, and decision-making^7–16^, constrains coding capacity. Finally, because primates perform visual recognition tasks nearly perfectly^1,2,5,6^, errors that could reveal the neural determinants of successful compared are rare.

Most work on the role of visual cortex in recognition memory has focused on late stage ventral stream areas like inferotemporal cortex (IT) and the human lateral occipital complex^2,17–19^. These studies emphasize average firing rates and linear decoding. More memorable images evoke stronger IT responses (“*magnitude coding”* ^2,17,19,20^) which is also observed in the late layers of artificial convolutional neural networks^20^. Familiar images evoke weaker responses (“*repetition suppression”; RS* ^2,17,21–25)^ (Figure 1, top). These findings suggest that behavior could reflect linear combinations of neuronal activity, a dominant view of how visual cortex encodes image features and identity^26,27^ and cognitive variables like attention, salience, and arousal or motivation^28–31^.

**Figure 1.**
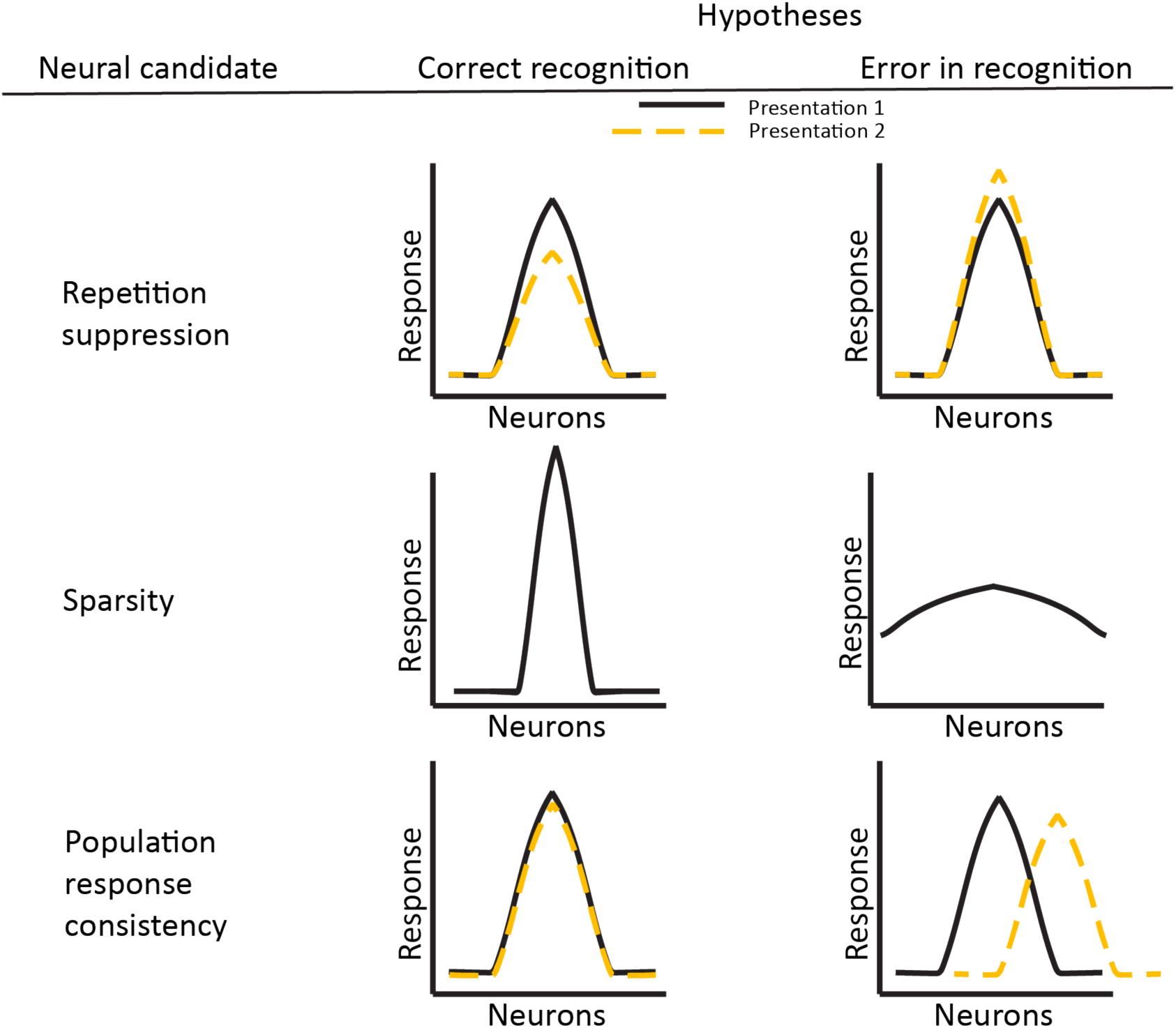
Three candidate, non-mutually exclusive neuronal signatures of successful recognition memory. The x-axis depicts the responses of visual neurons with varying tuning. Repetition suppression refers to a reduction in firing rate from presentation 1 to presentation 2, where stronger suppression could predict successful recognition. Sparsity refers to few neurons representing the stimulus. Efficient encoding could support proper storage necessary for successful recognition. Population consistency refers to the reliability of responses across presentations, such that more similar activity patterns could predict successful recognition. An extension of this idea is pattern completion, in which partial input can recapitulate neural activity that represents the entire stimulus. This would manifest as the second presentation requiring less image fragments to mimic the response profile of the first presentation.

Mid-level visual areas like V4 have been less studied but are well positioned to support memory-related computations. V4 encodes mid-level features linked to image memorability^32–35^, and reflects cognitive processes like attention, salience, learning, and working memory^16,36–40^. V4 responses to even static stimuli are dynamic^41^, raising the possibility that it could support temporally extended computations that have been implicated in memory^42–48^. Moreover, recognition memory distinguishes images that differ in only simple features and not semantic content^6,49^, implying that memory must rely on rich, feature-specific representations like those in mid-level visual cortex.

Here, we tested the hypothesis that V4 contributes to one-shot visual memory by evaluating which neural signatures distinguish between successful and unsuccessful memory (Figure 1). Inspired by perceptual decision-making studies that identify neural mechanisms by studying evidence accumulation^49^, we developed a visual recognition task for rhesus monkeys (Macaca mulatta) that temporally extends the recognition process. This design allowed us to dissect encoding and retrieval dynamics and compare neuronal population responses on correct and error trials. Monkeys viewed a natural image from an image set annotated for memorability and visual similarity^50,51^ and reported whether each was novel or had been seen once before while we recorded from populations of V4 neurons.

Our design enabled us to compare multiple candidate neural coding schemes including magnitude coding, repetition suppression (RS), trial-by-trial population response consistency, and dynamic pattern completion (Figure 1). Rate-based codes like RS^2,17,21,52,53^ are limited by trial-to-trial variability, which can blur responses to different images, inconsistent with primates’ near-perfect memory performance^6^. Other proposals, include *sparse coding* (Figure 1, middle), which could improve efficiency and reduce noise-related overlap in representations of similar stimuli^54–56^, as in pattern separation in the hippocampus^42,56,57^. Response consistency-based codes like *pattern similarity*^58–60^ and *pattern completion*^42–48,61,62^ (Figure 1, bottom), have not been evaluated in mid-level visual cortex. Because such codes have the capacity to support recognition memory, we hypothesized that reliable V4 population responses would be associated with successful memory.

Indeed, our V4 recordings reveal evidence of all proposed memory coding schemes, but the strongest predictors of correct recognition were population response consistency across first and second presentations and accelerated neuronal dynamics for familiar images, consistent with pattern completion. These signatures, traditionally associated with the hippocampus, align closely with behavior and outperform classic firing rate-based predictors.

Our results suggest that mid-level visual cortex contributes to high-capacity visual memory and that memory-related computations emerge earlier in the visual hierarchy than previously appreciated. By linking behavioral performance with population-level, trial-resolved analyses, we uncover coding mechanisms in V4 that may underlie fast, high-capacity learning and memory.

## Results

### A more difficult variant of a visual recognition memory task

Visual recognition memory is often studied in the context of a continuous recognition memory task^2,19,20,63,64^ (Figure 2a), in which animals view a series of many images and indicate whether each is ‘novel’ (never seen before) or ‘familiar (seen once before). The challenge with using this task to identify neuronal population signatures of successful memory is that memory is almost always successful^1,2,5,6^. This excellent performance precludes the opportunity to establish strong links between neurons and behavior by comparing neuronal responses on correct and error trials.

**Figure 2:**
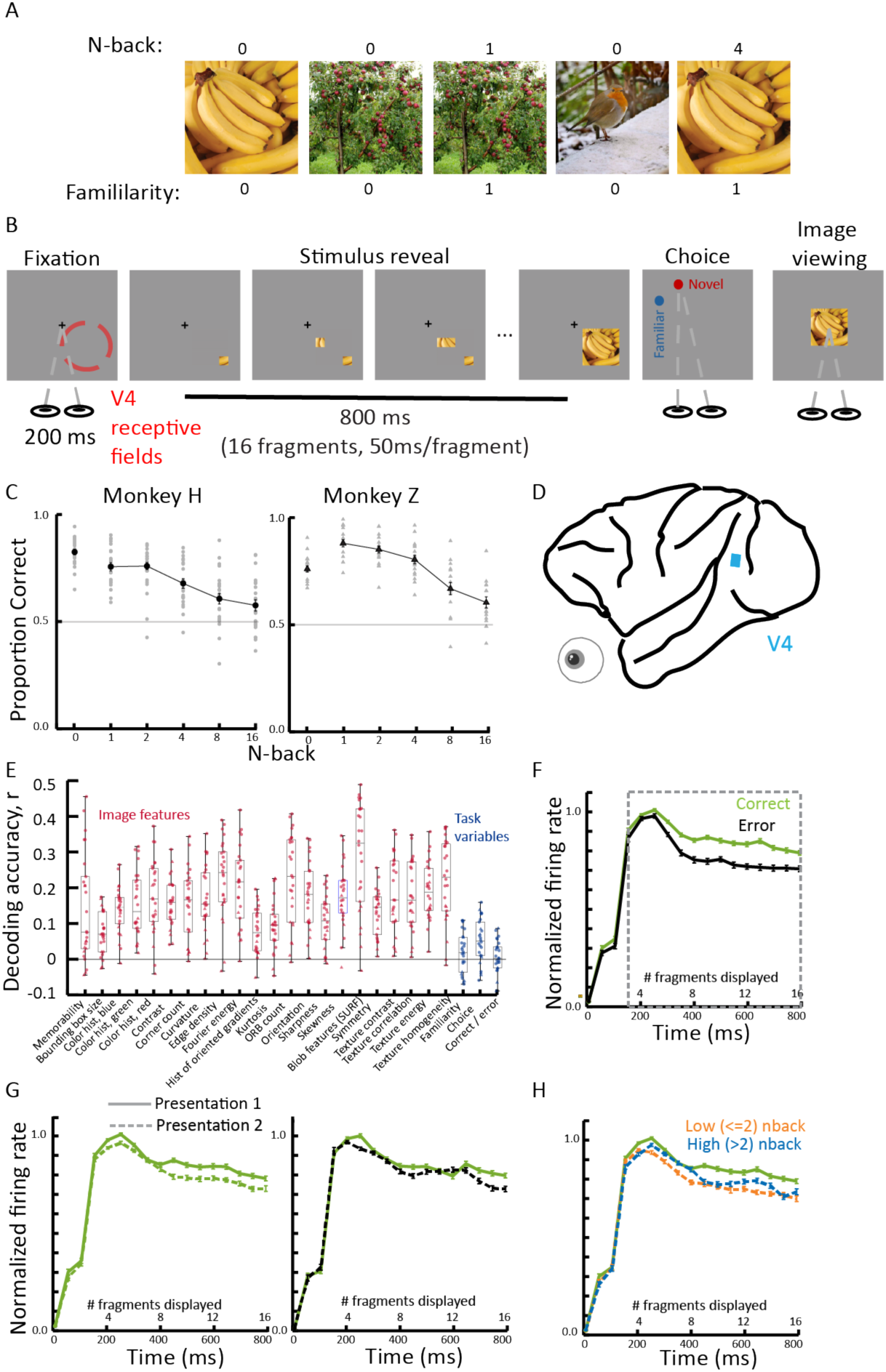
Task and recording methods, behavioral performance, and basic neural representations. **A: Depiction of the continuous recognition task.** Natural images are shown on each trial, and the subject makes judgments about whether the image has never been seen before (novel, familiarity = 0) or seen once before that session (familiar, familiarity = 1). The variable delays between the first and second presentations of an image are referred to as n-back. **B: Schematic illustration of the continuous recognition task used in our task.** The animal fixates to initiate the trial. The stimulus begins to appear one fragment at a time in the joint receptive fields until the entire image is displayed. After, the animal makes an eye movement according to his belief on whether the image is novel or familiar and is rewarded if correct. **C: Summary of behavioral performance for each animal.** The proportion correct is plotted as a function of n-back condition (0 = novel, 1:16 = familiar). Grey dots represent the performances of individual sessions, while black dots indicate the total average accuracy across sessions for each condition. Error bars denote the SEM. Performance across all n-back conditions was significantly above chance in both monkeys (p<.01). **D: Electrophysiological recordings.** We recorded extracellular spiking activity from neurons in V4. Data were obtained using chronic microelectrode arrays. **E: Linear decoding of image properties (red) and task variables (blue).** The correlation coefficient between the actual values and predicted values from our linear decoder was plotted for each variable. 0 represents chance. Each dot represents the average correlation coefficient for a single session. All image properties (red) had significant decoding accuracies (p<.01). For the task variables (blue): choice decoding was significantly better than chance (p = 0.0008). Image familiarity and correct vs. error trials were not significantly different from chance (p= 0.1513 and 0.4386, respectively). Statistical significance is assessed using a Wilcoxon signed-rank test against zero, testing whether decoding was above chance across sessions. **F-H: Peristimulus time histograms (PSTHs) for one example session, split according to image familiarity and behavior.** PSTHs were constructed by averaging and normalizing across all visually responsive units. Bins of 50ms were used, corresponding to the rate in which new fragments appear on screen. Line type denotes familiarity: trials of novel images consist of full lines, trials of familiar images consist of dashed lines. Error bars across all panels refer to SEM. Grey boxes illustrate the time period in which data were analyzed in E and all subsequent analyses. **F:** Displays the response to only first image presentations. The color of the line corresponds to if the animal was correct (green) or not (black). **G:** Displays the response to the first and second image presentations depending on the animals’ behavior. The color of the line corresponds to if the animal was correct (green) or not (black). Only trials in which the animal was correct for the first presentation were included. **H:** Displays the neural response to the first and second image presentations. For the second image presentation, the color refers to n-back condition (orange and blue correspond to trials of low or high n-backs, respectively). Only correct trials were included in this plot.

We overcame this challenge by developing a more difficult version of the continuous recognition task. On each trial, two rhesus monkeys (Macaca mulatta, both male, weights ∼13 kg) viewed a natural image^51^ that was gradually reviewed in a series of fragments (Figure 2b). They then indicated whether the image was novel or familiar by making an eye movement to the appropriate target. We manipulated image recency using an *n*-back design, with variable lags (1, 2, 4, 8, or 16 trials) between first and second image presentations (Figure 2a).

The gradual reveal slowed the decision process and increased task difficulty, enabling us to identify the components of the image, reveal dynamics, and neuronal population activity most strongly linked to behavioral success. Behavioral performance declined with increasing lag but remained above chance across conditions (Figure 2c). Across many sessions, monkey Z achieved an average accuracy of 78.0% (N = 16 sessions) and monkey H 74.3% (N = 22 sessions). Without the gradual reveal, animals performed near ceiling (Supplemental Figure S1), consistent with standard continuous recognition tasks. This drop in performance allowed us to compare neuronal activity on correct and error trials.

While the monkeys performed this task, we recorded spiking activity in V4 using a chronically implanted microelectrode array (Figure 2d; 64 channels). We chose V4 because of its well-established sensitivity to image features^34,35^ and its modulation by attention, learning, and memory^31^. In addition, the latency of image memorability^65^ and changes in human visual cortex following memory-boosting perturbations^66^ suggest an important role for mid-level visual cortex in visual recognition memory. Our experimental approach allowed us to test multiple proposed coding schemes, including magnitude coding, repetition suppression, sparse coding, population response consistency, and pattern completion, all within the same dataset.

### Image information can be decoded more successfully than task variables from populations of V4 neurons

Inspired by the widespread observation that V4 neurons are tuned for image features like orientation, color, and texture^34,35^, we and others have successfully used linear methods to decode image information from populations of V4 neurons. We replicated and extended these results here. We extracted 22 image features from the stimuli and used linear methods to decode each from the V4 population responses during the reveal period (Figure 2e; see Methods). Decoding performance varied across sessions but was significant for all image features (each red dot is one session, Figure 2e).

Surprisingly, we could also decode image memorability, which reflects the proportion of human observers that could successfully remember each image in standard versions of the continuous recognition task (quantified using scores from human psychophysical studies^50^). Memorability was linearly decodable from V4 (p = 0.0001), despite prior evidence that memorability sensitivity emerges in downstream areas such as inferotemporal cortex (IT) in monkeys^20^ and the lateral occipital complex (LOC) in humans^67^. Our results suggest that V4, which is an earlier stage of processing than those areas, contains linearly encoded signals related to memorability, perhaps due to the interplay of low-level features and memorability, and potentially species-specific differences. Interestingly, this finding is consistent with the result that memorability sensitivity emerges throughout the hierarchy of artificial neural networks^20^.

In contrast to image properties like features and memorability, task-related variables (blue dots, Figure 2e) were decoded less reliably. Choice decoding, our ability to predict the response (novel or familiar) of the monkey on the current trial, was the best, possibly reflecting premotor or decision-related signals. However, our ability to decode familiarity (presentation 1 vs. 2) and correct versus error trials were not distinguishable from chance, suggesting that the difference between successful and unsuccessful memory may not be captured by linear differences in average firing rates. These findings motivate a deeper look at alternative coding mechanisms beyond rate-based representations.

### V4 neurons respond more to novel than familiar images

We next replicated past observations that familiarity decreases responses of neurons in visual cortex, consistent with repetition suppression. We compared normalized population firing rates during the stimulus reveal on first and second presentations of each image, conditioned on behavioral outcome (correct or error). Each unit’s firing was scaled relative to its own mean, ensuring equal contribution to the population average. Unlike typical V4 responses to stimuli that fill the receptive field, the responses of our recorded units showed a delayed transient (150 ms to peak) and sustained activity, likely reflecting the gradual reveal of visual content within receptive fields (Figure 2f-h).

On first image presentations, responses were higher on trials with correct ‘novel’ judgments than on error trials, consistent with magnitude coding and potentially with attentional or arousal states in which mean firing rate might covary with performance^30^ (Figure 2f). Repetition suppression (the widely reported lower firing on second versus first presentations^2,17,21–25)^ was evident regardless of behavioral outcome, but was stronger when the animal successfully identified the image as familiar (Figure 2g). Suppression was also modulated by the number of trials between the first and second presentation (*n*-back): images repeated after shorter delays (≤2 trials) showed greater suppression than those with longer lags (>2), consistent with prior findings in IT^2^ (Figure 2h).

Together, these firing rate-based analyses demonstrate several properties of our task and the V4 neurons we recorded: the task is sufficiently difficult to allow us to compare neuronal signatures of correct and error trials, our V4 neurons robustly encode image features, including memorability, in ways that are decodable using linear methods, and these neurons show repetition suppression. These results put us in a good position to evaluate whether firing rate codes and other candidate neural mechanisms (outlined in Figure 1) can account for recognition memory capacity and performance.

### Repetition suppression does not predict successful memory when accounting for image recency

We first tested the hypothesis that repetition suppression (RS) in V4 could feasibly support recognition memory. This hypothesis predicts that there will be greater suppression to familiar images that are correctly remembered compared to those that were forgotten. Alternatively, repetition suppression might track image recency without contributing to memory. This hypothesis predicts more suppression for shorter n-backs (Figure 2a, 2h), but that the amount of suppression for trials with matched image recency does not relate to successful memory.

To tease apart these hypotheses, we quantified RS in individual units using a repetition suppression index (RSI) that is positive when responses to the second stimulus are less than the first (see methods). Most units had positive RSIs for both correct and error trials, consistent with broad suppression across the population (see Figure 3a for an example session), and there was greater suppression on correct than error trials on both this example session and across sessions for both monkeys (Figure 3b, Supplemental Figure S2). These results seem consistent with the idea that stronger RS is associated with successful memory.

**Figure 3:**
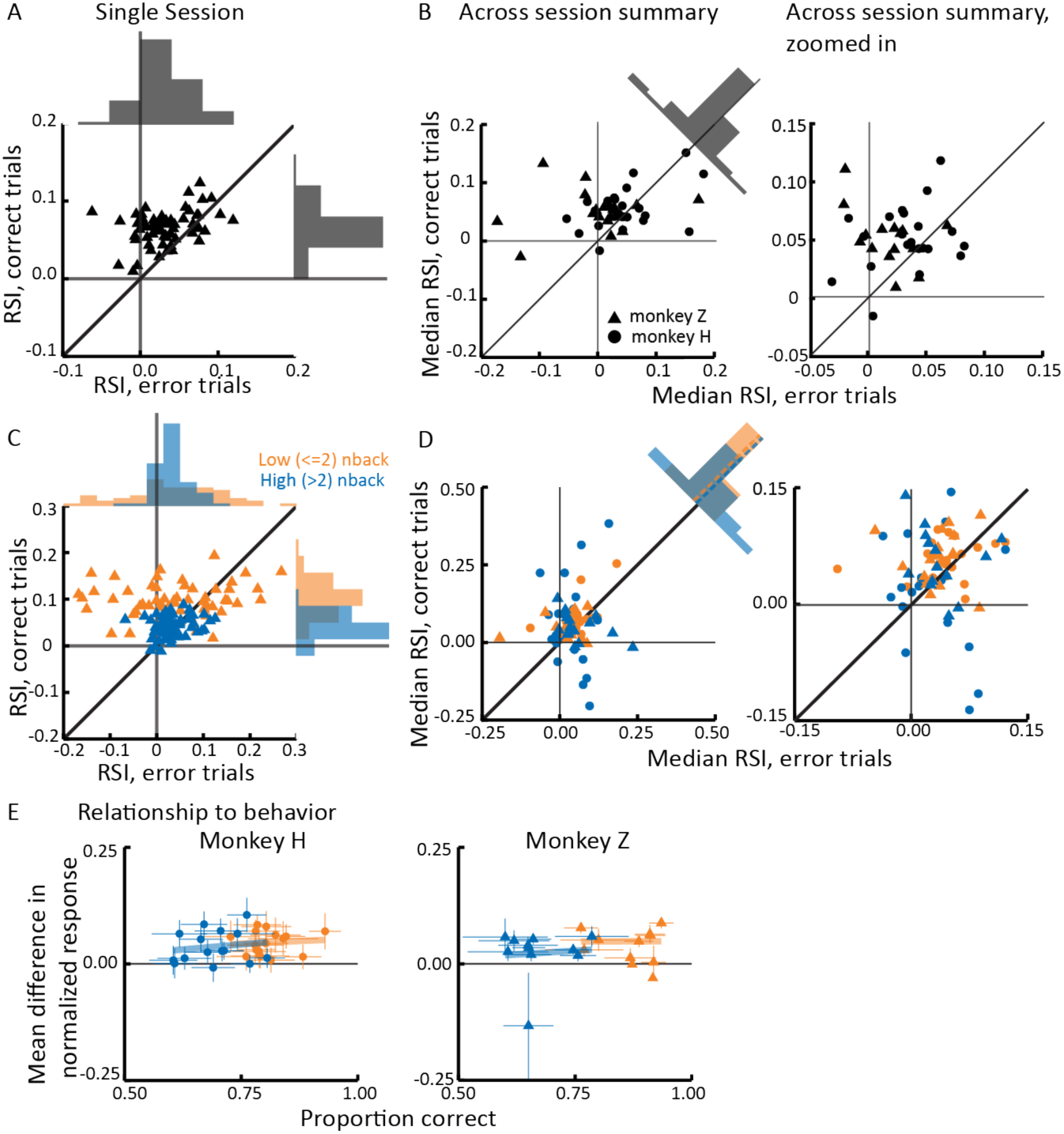
Repetition suppression does not predict successful memory when accounting for image recency. Dot type refers to the animal (circles = monkey H, triangles = monkey Z). Color refers to n-back condition: orange are low n-backs, blue are high n-backs, and black consists of all trial types regardless of n-back. **A: Single session example of repetition suppression indices (RSI) according to behavior.** The RSI (FR novel - FR familiar / FR novel) derived from each unit according to the animal’s behavior. In this panel, the RSI calculated from error trials is on the x axis and plotted against the RSI calculated from correct trials. Correct trials had larger RSIs than error trials (triangles cluster above unity line) (Wilcoxon signed-rank test, monkey Z, p=1.6181e-09). **B: Across session summary of RSI values according to behavior.** Comparison of the median RSI distribution from each session is plotted for correct versus error trials. The distribution is significantly skewed positive for correct trials (paired samples t test, both monkeys, p = 0.0046). **C: Single session example of RSI values considering n-back condition.** The same session as in panel A, but now each neuron represents two dots for both the n-back conditions. For each unit and n-back condition, we plot the average RSI on error trials (x-axis) against the average RSI on correct trials (y-axis). In the low back condition, correct trials had larger RSIs than error trials (Wilcoxon signed-rank test, p=1.24e-04), but not for high n-backs (Wilcoxon signed-rank test, p=0.0545). **D: Across session summary of RSI values considering n-back condition.** Similar to panel C, but now each session represents two dots for both the n-back conditions. Comparison of the median RSI distribution is plotted for correct versus error trials for each session and n-back condition. Significant differences were observed in the low n-back condition (paired samples t test, monkey Z: p = 0.0110, monkey H: p = 0.0235), but not in the high n-back condition in either monkey (paired samples t test, monkey Z: p = 0.6244, monkey H: p = 0.2607). Dashed lines in histogram indicate the median difference in RS between correct and error trials for each n-back condition. **E: Repetition suppression as a function of performance.** For each session, the normalized rate difference averaged across all units and images is plotted against the animal’s proportion correct. Error bars are SEM for each session. Line is fitted to the points using least squares method. Only sessions in which at least 250 valid image presentations (both trials across presentations completed) are included, resulting in 15 sessions from monkey H and 11 sessions from monkey Z (same sessions as Figure 4 panel F). Both low-back (monkey H: slope =0.04605, p= 0.98008; monkey Z: slope= −0.099468; p= 0.5109) and high n-back (monkey H: slope =0.086895, p=0.22389; monkey Z: slope= 0.064912; p=0.96698) had non-significant relationships with behavior (t-test, slope different than 0).

However, RS is also known to vary with the number of trials between the first and second presentations (*n*-back), in part because of its relationship to adaptation^68–73^. RS is stronger at shorter delays (Figure 2h), and behavioral performance is also better for low *n*-backs (Figure 2c). This means that correct trials are disproportionately drawn from low *n*-back conditions, potentially confounding RS with image recency. To disentangle these effects, we recomputed RSIs separately for low (*n* ≤ 2) and high (*n* > 2) *n*-backs. We chose this split to balance our ability to detect differences between low and high n-backs with the need to maintain enough trials for statistical analyses. In the example session (Figure 3c), correct trials showed stronger RS than error trials for low *n*-backs (orange points above diagonal), but this difference disappeared at high *n*-backs (blue points near diagonal). That is, at longer delays, we observed similar RS regardless of behavioral outcome.

We next looked at this effect across sessions. When we separated median RSIs by *n*-back condition (Figure 3d), we observed a small difference between RSIs on correct and error trials for lower n-back trials. This is not surprising because RS is very strong at small delays. However, the animals successfully perform this task at much larger delays, so a feasible neural mechanism must be consistent across delays. On larger n-back trials, RS was statistically indistinguishable between correct and error trials for both monkeys. Furthermore, there is was no significant relationship between mean RS across all units and performance on a session-by-session basis. These analyses suggest that RS is a robust marker of stimulus recency but is insufficient to support recognition memory on behaviorally relevant timescales.

### Unusually elevated activity in a sparse subset of neurons predicts successful memory

The use of natural images in this task allowed us to evaluate the hypothesis that sparse coding relates to successful memory. Sparse coding, in which a small number neurons are very active while the rest of the population is relatively inactive, has long been theorized to be efficient for coding image representations in visual cortex^54,55^ and memory representations in the hippocampus^42^.

Inspired by literature from the hippocampus, we hypothesized that visual memory in visual cortex could be supported by strong activation in a small subset of neurons during initial image exposure (i.e., the first image presentation). Some common measures of sparsity, like the skewness and kurtosis of firing rate distributions, were indistinguishable on correct and error trials (Supplemental Figure S3). We therefore tested whether rare, high-firing events in individual units predicted memory success. Figure 4a shows the distribution of normalized firing rates on an example correct and an example error trial. By definition, most units fired near their mean (z ≈ 0) in both trials, but the correct trial contained more units with z-scores greater than two (five units, compared to only one on the error trial). Across trials, the proportion of units whose activity was unusually high (Figure 4b) was greater on correct than error trials (Figure 4c). Across sessions, a larger proportion of units responded more than 2 standard deviations above their mean on correct than error trials (Figure 4d), even though this activity was by definition rare overall (typically <10% of units on any given trial, and different units contribute on different trials). Together, these findings support a sparse coding account in which successful recognition is associated with unusually elevated activity in a small, variable subset of V4 neurons during memory encoding.

**Figure 4:**
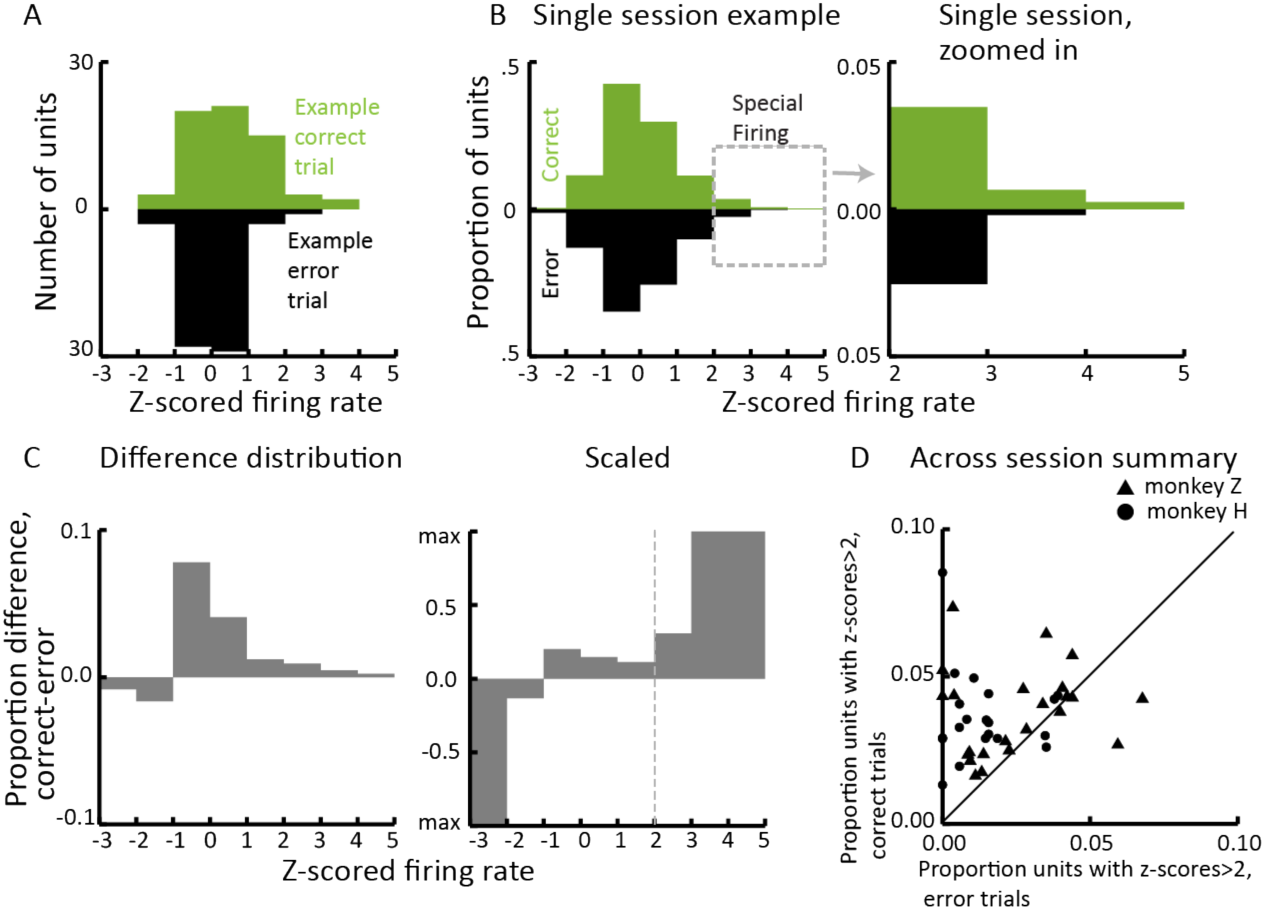
Unusually elevated activity in a sparse subset of neurons predicts successful memory on future image presentations. Example trial and session depicted in this figure is from monkey Z. All neural data comes from novel trials (first image presentation). In A and B, color corresponds to whether the animal was correct (green) or not (black) on familiar trials (second image presentation). Statistics use Wilcoxon rank-sum test. **A: Distribution of units’ z-scored firing rate in two example trials. B: Distribution of units’ z-scored firing rate for novel images in a single example session.** Left: The proportion of units within each z-scored firing bin for correct and error trials in the example session. Right: Zoomed in on the units with special firing. **C: Difference distribution between correct and error trials of the example session.** Left: Absolute difference distribution. Right: Difference distribution scaled by proportion of units. Similar to left but divided by the number of units in each bin. D: **Across-session analysis.** Each dot represents a session. The proportion of units above two standard deviations on correct trials (y axis) is plotted against the proportion on error trials (x axis). Distribution is significantly skewed larger for correct trials (monkey H, p = 0.00082308; monkey Z, p = 0.028196).

### Population response consistency predicts successful recognition

We next tested the hypothesis that successful memory relates to population response consistency. To this end, we calculated the correlation between the firing rates of all recorded units during the first and second presentation of an image (termed *population correlation*). Consistent with prior fMRI work on cortical reinstatement^58–60,66,74–77^, correct trials had significantly higher population correlations than error trials (Figure 5a,b; Supplementary Figure S4). This effect was robust across sessions, with nearly all sessions showing higher consistency on correct than error trials (Figure 5c).

**Figure 5:**
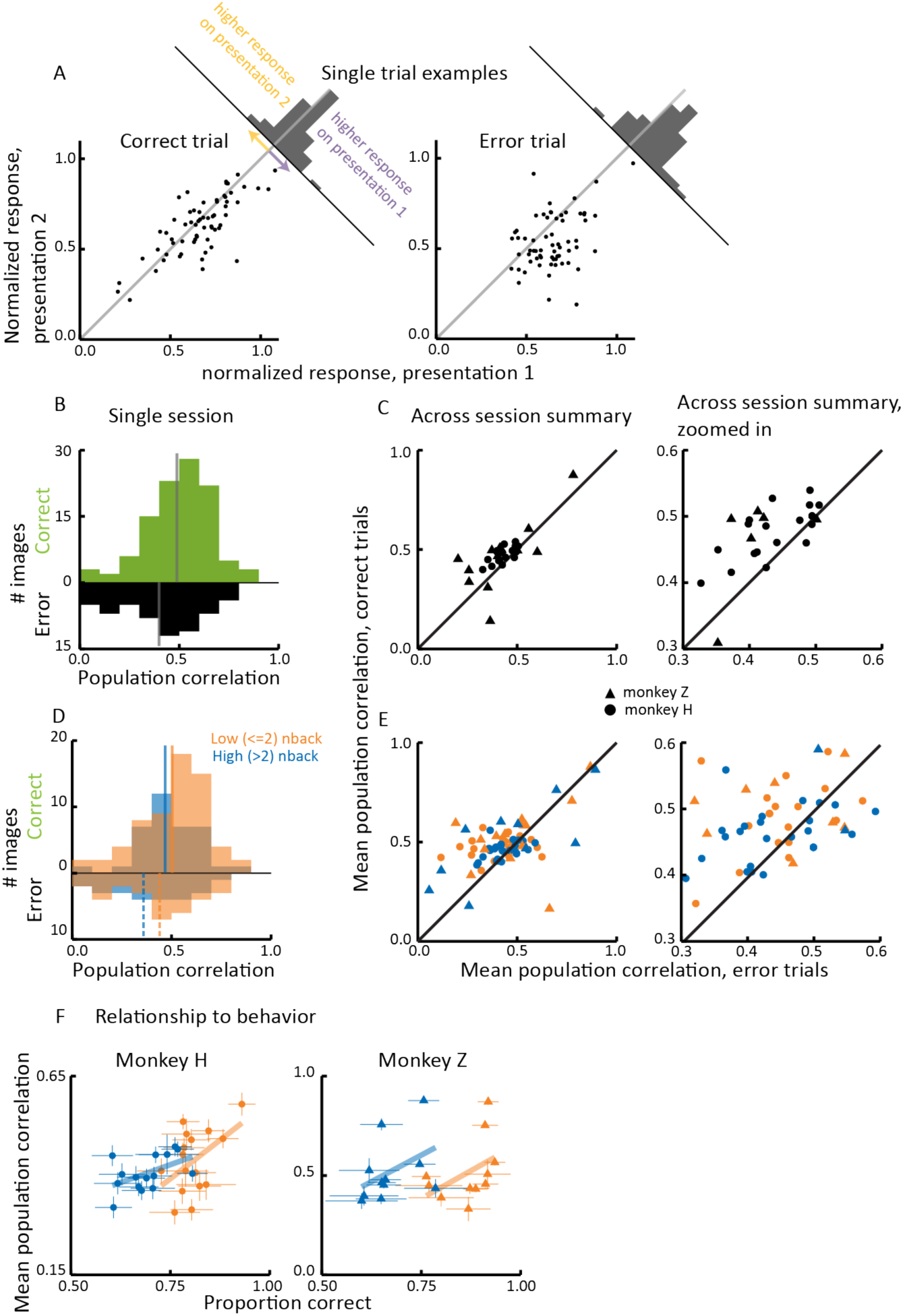
Population response consistency predicts successful recognition. Dot type refers to the animal (circles = monkey H, triangles = monkey Z). Color refers to n-back condition: orange points are low n-backs, blue are high n-backs, black consists of all trial types regardless of n-back. **A: Firing rates across presentations of two example images.** Each dot represents a unit’s normalized firing rate for presentation 1 (x axis) versus presentation 2 (y axis) of an image. Both presentations of the image were revealed in random, unique orders and were correctly identified as novel on the first presentation. Both examples show evidence of repetition suppression, with most points falling below the unity line. The left plot shows higher population response consistency (r = 0.757) and corresponds to a trial where the animal correctly identified the image as familiar. The right plot, with lower response consistency (r = 0.370), corresponds to an error in recognizing the image as familiar. **B: Spread of population correlation values for a single session of monkey H**. Distributions for familiar trials that were correct (green, top) versus errors (black, bottom) are displayed. Grey lines denote the median of the distribution. **C: Across session summary of population correlation values.** Comparison of the mean population correlation distribution from each session is plotted for correct versus error trials. The distribution is significantly skewed towards correct trials having larger values than error trials (paired samples t test, p=0.0008). **D: Spread of population correlation values for a single session considering n-back condition**. Similar to panel B, distributions for familiar trials that were correct (top) versus errors (bottom) are displayed. They are further split by low (orange) and high (blue) n-backs. The full and dashed lines indicate the mean in each n-back condition for correct and error trials, respectively. **E: Across session summary of population correlation values considering n-back condition.** Similar to panel C, but now each session represents two dots for both the n-back conditions. Comparison of the mean RSI distribution is plotted for correct versus error trials for each session and n-back condition. Both low (paired samples t test, p=0.0348) and high n-backs (paired samples t test, p=0.0145) have significantly larger population correlations for correct trials. **F: Population response consistency as a function of performance.** For each session, the average population correlation across all units and images is plotted against the animal’s proportion correct. Error bars are SEM for each session. Line is fitted to the points using least squares method. Only sessions in which at least 250 images in which both trial presentations were completed are included, resulting in 15 sessions from monkey H and 11 sessions from monkey Z (same sessions as Figure 2 panel E). Low n-back (monkey H: slope=0.77824, p=0.0021217; monkey Z: slope = 1.1231, p=0.024422) and high n-back (monkey H: slope=0.35964, p= 0.0.026846; monkey Z: slope =1.0662, p=0.010675) had generally consistent slopes, with all conditions exhibiting a significant relationship with behavior (t-test slope different than 0).

Because low *n*-back trials have shorter delays and are easier (Figure 5c), correct trials are more likely to come from these trials, which presents a potential confound. We therefore calculated population correlations separately for images with low (<=2) and high (>2) n-back. Correct trials maintained larger population correlations in both n-back conditions (Figure 5d; Supplemental Figure S4). This consistency difference between correct and error trials was present in nearly all sessions (Figure 5e), indicating that it is not driven solely by delay length or task difficulty.

In Monkey H, population correlations were higher on low *n*-back trials, (Wilcoxon signed rank test, p=0.0092) whereas Monkey Z showed no significant difference (Wilcoxon signed rank test, p= 0.9176). This likely reflects difference in the self-initiated inter-trial timing (Monkey H mean 7.4263 s; Monkey Z mean 4.5673 s).

Population response consistency also predicted overall performance during each session. The mean population correlation correlated strongly and significantly with behavioral performance at both low and high *n*-backs and in both monkeys (Figure 5f).

Together, these findings demonstrate that consistent population activity across image presentations is a robust predictor of successful recognition and memory retrieval.

### Temporal dynamics of population response consistency reveal evidence of pattern completion

Our task design allowed us to examine the temporal evolution of population response consistency and its relationship to successful memory. Pattern completion, which has been widely studied in the hippocampus^42,43,57,78^ but not in visual cortex, occurs when a partial sensory input triggers reinstatement of a full memory. We tested whether a similar phenomenon occurs in mid-level visual cortex.

Time-resolved population response consistency revealed higher time-by-time correlations for correctly than incorrectly recognized images (Figure 6a, b). This difference was especially notable along and below the diagonal, indicating that population responses to correctly recognized images became more consistent across time and stabilized more quickly. Antisymmetric difference matrices reveal positive values to the right of the diagonal on correct trials, consistent with accelerated responses, and zero or negative values on error trials, consistent with slower dynamics (Figure 6c,d).

**Figure 6:**
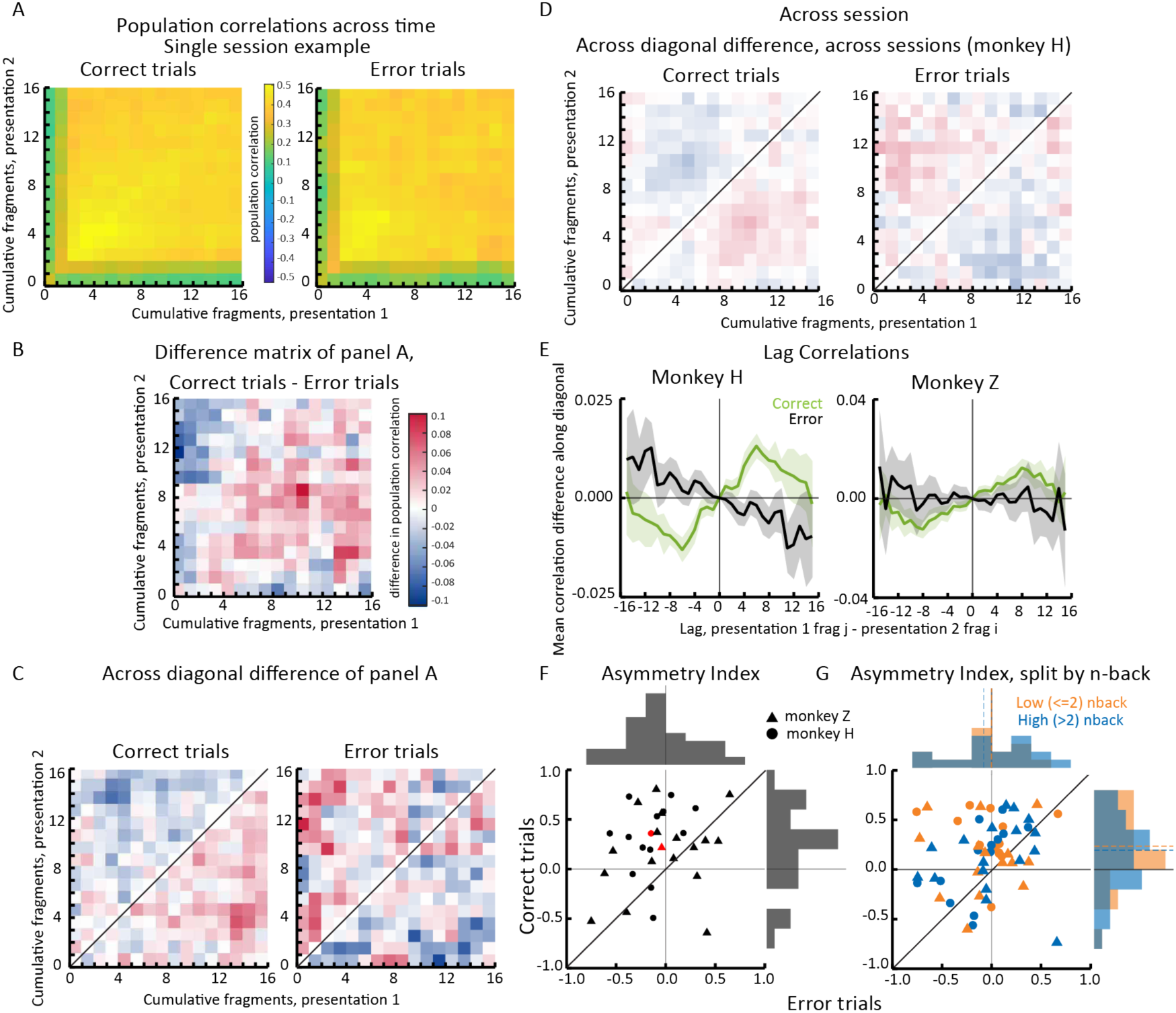
Temporal dynamics of population response consistency reveal evidence of pattern completion. **A: Single session example of population correlations throughout the stimulus reveal**. Each time bin corresponds to one image fragment. **B: The difference between the correct and error matrices from panel A.** **C: Difference across diagonal of matrices in panel A.** Left corresponds to the correct trials, right corresponds to the error trials. Same scale bar as panel B. **D: Across session summary of difference across diagonals.** Sessions from monkey H. Left corresponds to the correct trials, right corresponds to the error trials. Same scale bar as panel B. **E: Lag correlation for correct (green) and error (black) trials, averaged across all sessions**. Left is monkey H, right is monkey Z. Shaded region corresponds to SEM. Positive lags on the x axis represent earlier time points in presentation 2 and later time points in presentation 1. Negative lags represent later time points in presentation 2 and earlier time points in presentation 1. **F: Per session summary of pattern completion via the asymmetry index.** The average ASI values derived from the matrix of the across-diagonal difference of the population correlation matrix (panel C) for each session. The indices calculated for error trials from a session (x axis) are plotted against the indices for correct trials of that session (y axis). The black dots represent individual sessions and consist of all trial types regardless of n-back. Dot type refers to the animal (circles = monkey H, triangles = monkey Z) in this panel and in panel D. Red dots demonstrate the median index for each monkey. Correct trials in monkey H were significantly larger than error trials (paired samples t test, p=7.4207e-04), and correct trials had indices that were significantly skewed positive (1 sided t test, 0.0059943). Error trials did not; they were hedged negative (1 sided t test, p=0.025096). Correct trials in monkey Z were in the same direction but not significantly larger than error trials (paired samples t test, p=0.081979). **G: Per session summary of pattern completion via the asymmetry index considering n-back condition.** Color refers to n-back condition: orange are low n-backs, blue are high n-backs. Dot type refers to animal identity as in panel D. The orange and blue dashed lines in the histograms demonstrate the median index in each respective n-back condition. Low n-backs trended towards having larger positive indices than high n-backs (paired samples t test, p=0.0835). Correct trials across monkeys were significantly larger than error trials in both n-back conditions (paired samples t test; low n-back: p = 0.0048, high n-back: p= 0.0236), and were significantly positive (1 sided t test, low n-back: p =6.3761e-04, high n-back: p= 0.0408).

Lag correlation analyses confirmed this acceleration (Figure 6e). For correct trials, peak similarity occurred when the second presentation was shifted earlier relative to the first, consistent with a faster unfolding of the response. The peak lag occurred after a few hundred ms (six time bins for Monkey H, eight time bins for monkey Z). Error trials showed no shift or delayed responses.

This result was not explained by stimulus differences across image presentations. In control sessions in which sometimes the image was revealed in the same sequence across presentations, correct trials still showed positive lag correlations (Supplemental Figure S5), confirming that faster reactivation reflected neuronal population dynamics rather than stimulus details.

We quantified these faster dynamics on second presentations using an asymmetry index (Figure 6f). Both monkeys had positive indices (faster dynamics for second than first presentations). Monkey H showed significantly faster dynamics on correct than error trials. Monkey Z, showed a similar but not significant trend. Pattern completion was strongest for low n-back trials but remained significant for correct trials regardless of n-back condition (Figure 6g).

These findings indicate that during successful recognition, V4 population activity during the second image presentation advances more quickly along a familiar trajectory. This temporal shift is consistent with pattern completion, a hallmark of hippocampal memory retrieval^42,57,78^, now demonstrated in mid-level visual cortex.

Together, our results suggest that visual recognition memory is supported not just in firing rate-based codes, but also by the reliability and temporal evolution of population-level representations. Partial correlation analyses revealed that repetition suppression, sparse coding, and magnitude coding are interrelated through their shared dependence on overall firing strength (Supplementary Figure S6). In contrast, population response consistency, the variable most strongly linked to behavior, was largely independent of firing rate-based measures. These findings highlight the importance of population response reliability and dynamics in supporting visual recognition memory and potentially other high-capacity functions.

## Discussion

Our findings unify distinct neural mechanisms proposed to underlie visual recognition memory and highlight an important role for mid-level visual cortex in supporting high capacity. During memory encoding (presentation 1), high firing in a small subset of neurons, consistent with sparse coding, predicted later recognition. During retrieval (presentation 2), population response consistency was the best predictor of success: correct trials showed higher correlations between population activity patterns than errors. This consistency unfolded over time as accelerated neuronal dynamics, consistent with pattern completion in which population representations were reactivated more quickly on correct than error trials. Repetition suppression, although consistently present, primarily reflected image recency rather than behavioral success. These results identify sparse coding during encoding and temporally structured population consistency during retrieval as key neural signatures of successful visual memory in mid-level visual cortex.

### The role of response consistency in visual cognitive behaviors

Population response consistency has been linked to explicit visual memory in fMRI studies of visual and higher-order areas^58–60,79^. We show for the first time that V4 spiking activity exhibits this relationship. In contrast, repetition suppression^2,17,21^, while observable, was related to image recency and did not consistently distinguish correct from error trials, consistent with findings that it persists even under anesthesia^80–85^. Unlike repetition suppression, population consistency depends on the visual content of an image, which highlights its relevance to visual memory, which depends strongly on image properties^32,33,50,67^.

Our approach quantifies population-level consistency in a way that is distinct from previous work on noise correlations (correlations in the trial-to-trial fluctuations of two neurons across many repetitions of the same stimulus^7^), which have been linked to many visual cognitive processes^8–16^, and representational drift, which quantifies the long-term instability of single-neuron responses without behavioral change^86–90^. Although all of these metrics reflect neuronal response reliability, our approach can be measured even on pairs of trials: we measure the correlation between two trials across many neurons. This provides the opportunity to link response consistency to behavior under more naturalistic conditions, which lack extensive repetition of ostensibly identical conditions. Together, our findings along with literature on noise correlations and representational drift underscore the importance of neural reliability in cognitive function.

### Pattern completion in mid-level visual cortex

Pattern completion, typically associated with the hippocampus^43,91^, was evident in V4. The extent and accuracy of pattern completion correlated with memory performance (Figure 6). Unlike previous studies using artificial (optogenetic) reactivation in mice^92,93^, we observed spontaneous pattern completion during goal-directed behavior. Complementary fMRI ^94^ and inferotemporal cortex physiology studies suggest similar cortical effects^95,96^, potentially arising from feedforward^95^ or recurrent processing within visual areas^96^ or feedback from memory systems such as the hippocampus^66,74,97^.

Although our dynamic stimulus design reveals temporal signatures of pattern completion, the decision process remains unresolved. It is unclear whether animals integrate visual fragments until a decision threshold is reached or whether a single diagnostic fragment can trigger an immediate decision, akin to evidence accumulation studies using a drift diffusion framework^49,98–103^. Future tasks could determine how particularly memorable fragments influence accumulation.

### The role of mid-level visual cortex in visual memory

We demonstrate that memory-related signals emerge in mid-level visual cortex, which is earlier in the processing hierarchy than previously appreciated. Our data suggest a multiplexed code in V4, where image-related representations rely on a linear code (Figure 2) and memories use a code related to sparseness and response consistency. This multiplexed coding could support behavioral flexibility by allowing downstream areas to access different representations depending on task demands^104–106^ and may increase coding capacity, allowing the same population to support multiple functions simultaneously.

These memory signals are likely not exclusive to V4. Human fMRI studies show similar response consistency across occipital, parietal, and medial temporal areas during visual memory tasks^58–60,79,107^. Our findings are consistent with theoretical models proposing a division of labor, with hippocampus supporting episodic memory and cortex storing memory content ^48,78,108^. Evidence from human LOC^58,59^, which is often considered analogous to macaque IT and shares properties with macaque V4, supports widespread cortical involvement in visual memory ^109–111^. A recent study TMS study showed memory-related activity in the middle occipital gyrus^66^, reinforcing the idea of distributed visual memory representations.

### Opportunities for future study

Our task design, with gradual stimulus reveal, increases difficulty and isolates neuronal signals that distinguish correct from error trials. It enables manipulation of fragment order, content, and proportion of the image revealed, facilitating tests of visual properties that support memory, changes in disease, and temporal unfolding of recognition.

A major open question is how population response consistency might be read out to guide recognition on individual trials. Response consistency is ultimately a similarity metric between two trials and figuring out which memory to retrieve and compare poses a significant challenge to readout mechanisms. The key may be understanding the signals related to decision-making in memory tasks, perhaps from association areas in parietal and frontal cortex.

Together, our findings demonstrate that mid-level visual cortex encodes signals that differentiate success from failure in memory encoding and retrieval. Sparse activity predicts successful encoding, while consistent, temporally accelerated population responses predict successful retrieval. These mechanisms emerge after a single exposure to complex natural stimuli, suggesting that visual cortex plays a more active role in memory than previously appreciated. By dissociating behaviorally relevant neuronal signals from those that reflect stimulus recency, this work highlights the value of trial-by-trial analyses in naturalistic tasks. As neuroscience seeks to explain the brain’s capacity for flexible, rapid learning, and artificial intelligence strives to replicate it, these findings offer a powerful model of how high capacity, one-shot learning can emerge from sensory population codes.

## Materials and Methods

### Subjects and Surgery

We studied two adult male rhesus macaques (monkey H and Z, Macaca Mulatta; weight, 12–15 kg). Monkey H was experimentally naive; monkey Z had been used in previous tasks. All animal procedures were approved by the Institutional Animal Care and Use Committees of the University of Chicago, and all training, surgery, and experimentation methods were performed in accordance with the relevant guidelines and regulations. Standard aseptic surgical procedures under gas anesthesia were performed to implant a titanium head restraint device before behavioral training. Once training was complete, we implanted a 8 x 8 multielectrode arrays (Blackrock) in the left hemisphere of cortical area V4. We identified area V4 with stereotactic coordinates and by visually inspecting the sulci. We placed the array between the lunate and the superior temporal sulci.

### Experimental apparatus and visual stimuli

All visual stimuli were displayed on a linearized CRT monitor (1,024 × 768 pixels, 120-Hz refresh rate) placed ∼57 cm from the animal. We monitored eye position using an infrared eye tracker (Eyelink 1000, SR Research) and used custom software (written in Matlab using the Psychophysics Toolbox^112^ to present stimuli and monitor behavior. We recorded eye position and pupil diameter (1,000 samples per s), neuronal responses (30,000 samples per s) and the signal from a photodiode to align neuronal responses to stimulus presentation times (30,000 samples per s) using hardware from Ripple. Visual stimuli consisted of natural images with verified metrics (image labels, memorability scores, similarity scores, etc) from the THINGS^51^ dataset. During recording, all images used had never been seen prior to recording except for a subset of five images that were shown every day, termed the super familiar images. We positioned the images during the stimulus reveal to be centered and fit to the joint receptive fields of our V4 units (see final paragraph in Electrophysiology).

### Behavioral Task

We created a variant of a classic paradigm used to test visual recognition memory, the continuous recognition task. The nature of the task was for subjects to view images and to determine whether each image was novel (never before seen) or familiar (seen exactly once before). The image was picked randomly, either from the images that have never been seen before or a familiar image that was in one of the n-back delays, or an image that had been seen across session (super familiar, not analyzed in this paper). Throughout the session, on each trial there was an equal chance of the image picked being novel or familiar. During the task, monkeys fixated in a central fixation point (within a 0.5° fixation window) for 200 ms before the stimulus reveal period. During the stimulus reveal, fragments of the image were displayed in series (50 ms each, 16 fragments total) in a random order. Images were square and thus all fragments were equal sizes. Once an image fragment was on, it remained on screen until all fragments (i.e., the entire image) were shown. After the stimulus reveal, the image and fixation dot disappeared. This served as the go cue for the animals to report their choice on whether the image was novel or familiar by looking at one of two peripheral targets. To choose a target the animal was required to fixate on it for approximately 500ms. If the correct target was chosen, the animal was rewarded with a liquid reward. All trials in which the monkeys successfully completed the trial (rewarded or unrewarded) were analyzed. If the animal left the fixation window at any time during the stimulus reveal, the trial was aborted and data were discarded. After either a choice or trial abortion, the animal was required to fixate on the image for 500ms before moving on to the next trial. The average time to complete a trial was approximately 3.3s, and the average number of completed trials in one session was 472 for monkey H, 869 for monkey Z.

### Electrophysiology

Extracellular recordings were obtained in both monkeys using chronically implanted V4 electrode arrays. The array was connected to a percutaneous connector that allowed daily electrophysiological recordings. The distance between adjacent electrodes was 400 μm, and each electrode was 1 mm long.

We defined analyzable recording sessions as those with at least 250 completed trials in a session (completed meaning the subjects successfully maintained fixation until they indicated their choice), unless otherwise stated. This criterion resulted in 22 recording sessions for monkey H and 16 sessions for monkey Z. We amplified and filtered the neural activity by setting the threshold for each channel at three times the standard deviation and used threshold crossings as the activity on that unit. We recorded a combination of single neuron and multiunit clusters. We use the term “unit” to refer to either. Analyses comparing single and multiunits in previous work^8,113^, did not find systematic differences between single neurons and multiunits for the types of population analyses presented here. For that reason, we analyzed multiunit activity (one multiunit per channel).

To determine which units to include from each recording session, we required that the transient response to the stimulus was at least 10% more than baseline activity (measured in the 200 ms fixation period preceding the stimulus reveal period). Units that did not reach this criterion were excluded from further analysis. The average number of included units per recording session was 64 for each monkey. In order to control for artifact noise that may be shared across multiple channels, we excluded trials in which the activity of more than 50% of units was greater than three standard deviations away from the mean activity for that unit.

The receptive fields of these visually responsive units were measured using separate mapping sessions. The monkeys fixated their gaze on a dot while Gabor stimuli were subsequently flashed at random locations for 100ms each across the screen. The receptive fields of our units were contralateral to the array site and well contained within the boundaries of our display screen. They were approximately 5° eccentric and a similar width size (a property typical for V4 receptive fields). We did not observe considerable changes in receptive field location over the duration of the recording period.

### Data Analyses

Analyses of spike data and statistical tests were performed using custom software written in MATLAB (MathWorks). All analyses were performed using spikes from the stimulus reveal from 150ms to 800ms (window outlined in Figure 2f), unless otherwise stated. This window was chosen by considering the typical V4 latency along with the slower dynamics due to our task design. Only trials where both trials of image presentations were completed were included.

#### Linear Decoding

We used cross-validated linear decoding methods to determine our ability to differentiate neuronal population responses between two categories. All decoders were determined separately for each animal, decoded feature, and recording session, and utilized spike counts (averaged over the analysis window of the stimulus reveal period for decoding performances) (Figure 2f). The decoder was a linear classifier trained to best differentiate the V4 population responses to certain image features (such as memorability) or task variables (such as choice). For each session, we performed 10 random train/test splits, using half the trials for training and the other half for testing. In each split, a linear model was fit to the training data, and decoding performance was quantified as the Pearson correlation coefficient between predicted and actual labels on the held-out test data. The final decoding accuracy was the average correlation across the 10 repetitions.

We extracted features for all images as follows. Memorability scores were publicly available for the THINGS image set; scores were derived from human behavioral data^50,51^. All other image features were extracted using custom MATLAB code: each image was first rescaled to 800×800 pixels and then converted to grayscale before being passed to a variety of built-in MATLAB functions (MathWorks 2024b). The list of image features and their method of calculation is as follows: bounding box size (normalized area of the smallest bounding box around the largest object in the binarized image), color histogram blue (average intensity of blue pixel values, normalized per image), color histogram green (average intensity of green pixel values, normalized per image), color histogram red (average intensity of red pixel values, normalized per image), contrast (standard deviation), corner count (total number of corners calculated from a Harris corner detector), curvature (standard deviation of the Laplacian transform), edge density (average fraction of edge pixels calculated from a Sobel edge detector), fourier energy (average fast fourier transform), H.O.G. (average histogram of oriented gradients calculated from detecting local shape information in an 8×8 pixel window), kurtosis (tailedness of distribution of pixels), orb count (number of keypoints calculated from an Oriented FAST and rotated BRIEF feature detection method), orientation (average vector of gradient directions), sharpness (variance of the Laplacian transform), skewness (asymmetry of distribution of pixels), blob features (SURF) (number of blob features calculated from a Speeded-Up Robust Features algorithm), symmetry (difference in normalized pixel intensity between image and mirrored version of itself), texture contrast (contrast statistic extracted from normalized gray level co-occurrence matrix), texture correlation (correlation statistic extracted from normalized gray level co-occurrence matrix), texture energy (energy statistic extracted from normalized gray level co-occurrence matrix), texture homogeneity (homogeneity statistic extracted from normalized gray level co-occurrence matrix). The list of task variables and their definitions are as follows: familiarity (whether the image was presented for the first or second time), choice (saccade target), correct / error (whether the chosen saccade target was correct).

#### Repetition Suppression Index

To quantify the amount of repetition suppression in each unit (Figure 3b), a repetition suppression index (RSI) was computed, as follows:

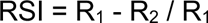

Where R1 and R2 refer to the responses on the first and second image presentation. This index ranges from −1 to 1 to quantify the strength and directionality of differences in spike rates across images. A negative index represents repetition enhancement (the second presentation elicited higher rates compared to the first presentation), whereas positive represents repetition suppression (the second presentation elicited lower rates compared to the first presentation). Only trials in which trials of both image presentations were completed were used.

#### Asymmetry Index

To quantify the amount of pattern completion occurring on a trial, an asymmetry index (ASI) was computed on the difference matrix to better portray the imbalance of time dynamics on presentation 1 versus presentation 2. The difference matrix was calculated from the population correlation matrix across time (M) as follows:

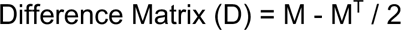

This isolates the antisymmetric component of D, where Dij=−Dji. We then compute the Asymmetry Index (ASI) as follows:

Define:

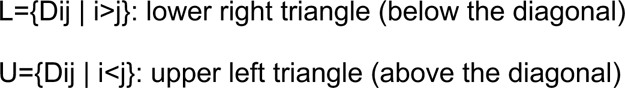

Then:

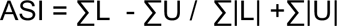

This index reflects the directional imbalance between the lower and upper triangle of the antisymmetric component, normalized by total antisymmetric magnitude. Thus, these values are locked between −1 and 1, with a positive index indicating more weight in the lower triangle (suggesting faster activity during the second presentation). Negative index values indicate the opposite: slower activity during the second presentation. Only trials in which both the first and second image presentations were completed were used.

#### Lag correlation

To understand the temporal structure of response consistency, we computed a lag correlation. This was calculated separately for correct and error trials using the difference matrix (Figure 6d, defined above). A lag correlation was then computed by averaging the values along each lag (ie., matrix off-diagonal) of D, corresponding to a specific temporal lag between the two image presentations. Positive lags represent earlier time points in presentation 2 and later time points in presentation 1; negative lags represent later time points in presentation 2 and earlier time points in presentation 1. Shaded error bands reflect the standard error of the mean (SEM) across sessions. Only trials in which both the first and second image presentations were completed were included in the analysis. This approach allows us to assess whether activity patterns during the second image presentation are temporally shifted relative to those during the first, and whether this shift differs between correct and error trials.

## Resource availability

Data and code are available from the lead contact upon request.

## Acknowledgements

We are grateful to Kursti Eaves for providing training and recording assistance, and to Ramanujan Srinath for his contribution to the linear decoding analysis. We would also like to thank Douglas Ruff, Wilma Bainbridge, and Joel Voss for their help in improving the manuscript. This work is supported by the Simons Foundation (Simons Collaboration on the Global Brain award 542961SPI) and the National Institutes of Health (awards R01EY022930, R01EY034723, RF1NS121913, and 5T90DA059109-03).

## Author Contributions

Conceptualization – GFD, CX, MRC

Data collection – GFD, CX

Experimental software – GFD, CX

Formal analyses – GFD, MRC

Writing, original draft – GFD, MRC

Writing, review & editing – GFD, CX, MRC

Funding acquisition – MRC

## Supplemental Information

**Figure S1:**
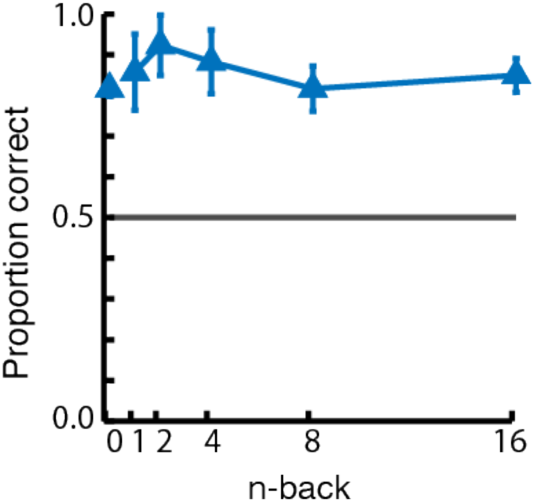
Behavioral performance on the continuous recognition task without the gradual reveal. Average accuracy across n-back was computed across three sessions in monkey Z. Error bars show SEM. Performance is better across all n-backs compared to the performance with the gradual reveal in Figure 2c.

**Figure S2:**
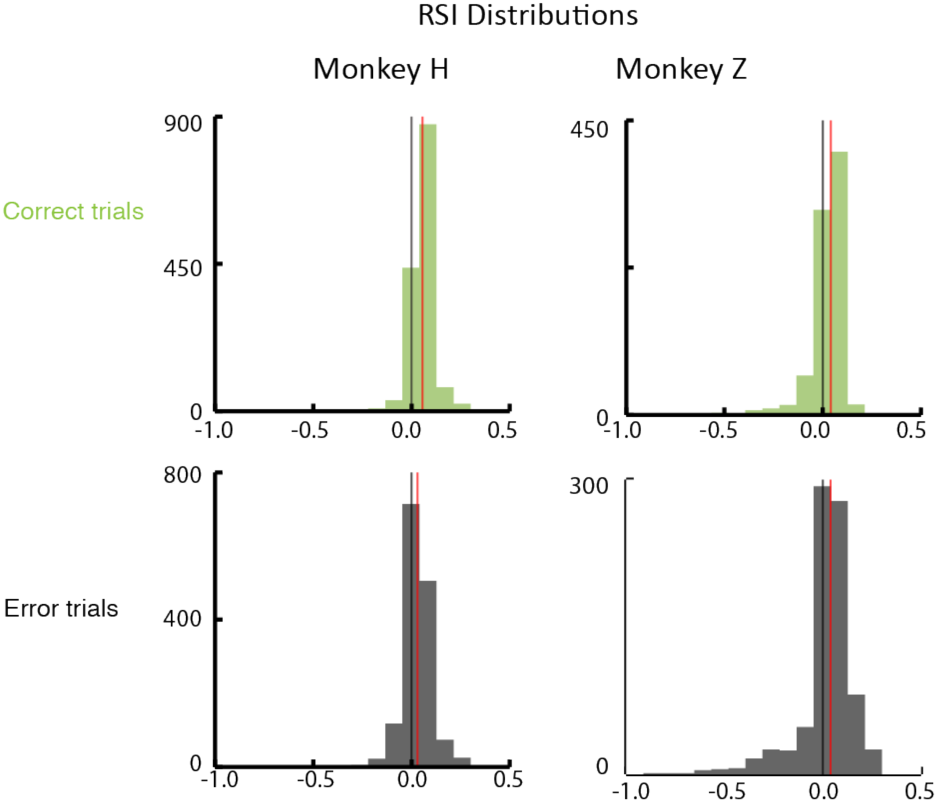
RSI distributions, split by monkey and behavior. Columns differentiate monkeys, rows differentiate whether the animal correctly recognized the image or not. Red vertical lines represent median RSI for the respective condition. Black vertical lines mark zero. For every session, we plot the average RSI value derived from each unit (see methods). All distributions were shifted towards positive values, indicating repetition suppression in all monkeys and conditions. Monkey H had significantly more repetition suppression on correct trials compared to errors (rank sum test, 20 sessions,p= 9.1150e-45). Monkey Z did not (rank sum test, 13 sessions, p=0.4655).

**Figure S3:**
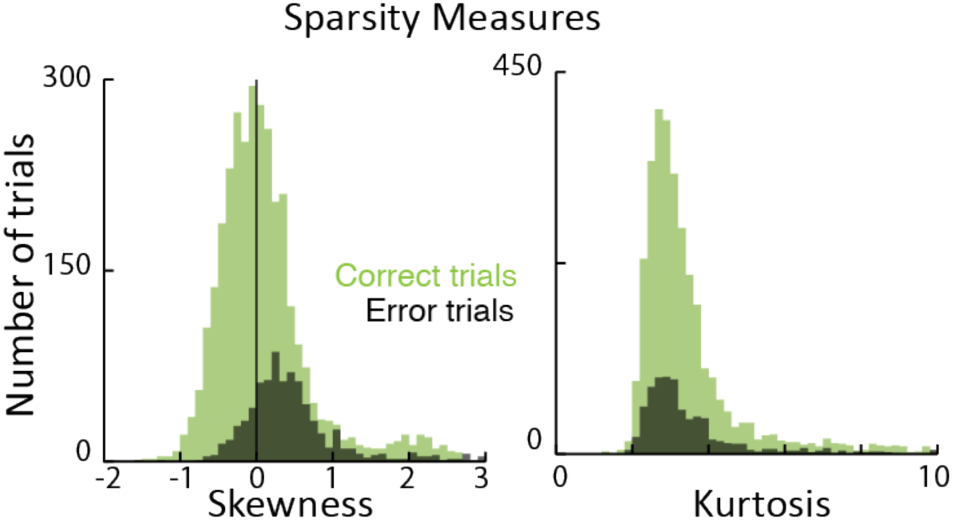
Sparsity measures. Skewness (left) and kurtosis (right) of z-scored firing rate distributions on first image presentations. Color refers to whether the animal was correct (green) or made an error (black) on the next image presentation. No significant difference in the skewness (Wilcoxon rank-sum test, p = 0.9935) or kurtosis (Wilcoxon rank-sum test, p= 0.78957) for correct versus error trials.

**Figure S4:**
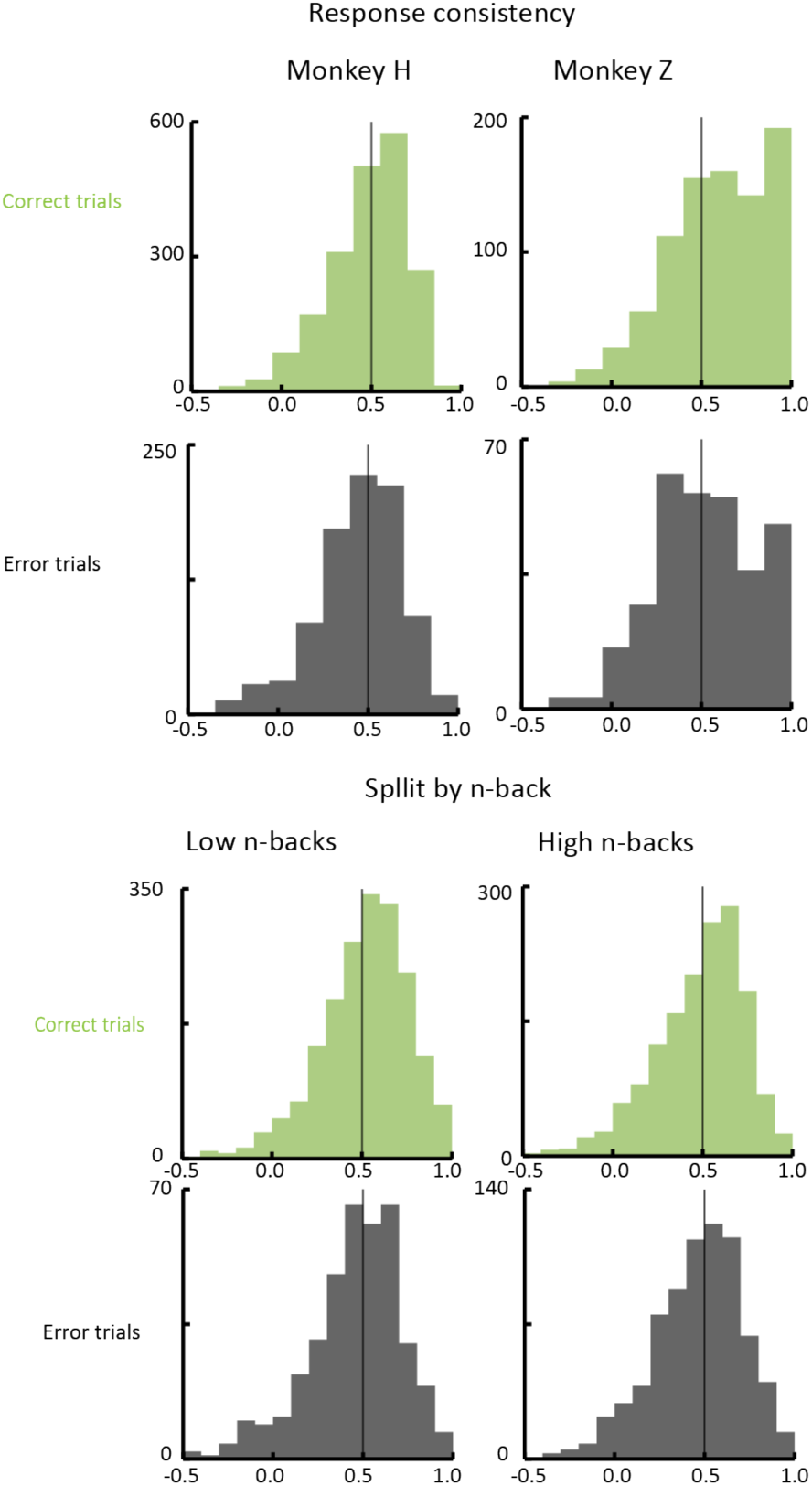
Population correlation distributions. Top shows differences in the distributions for each monkey (column) according to whether the animal correctly recognized the image or not on the second presentation (rows; green is correct, black is errors). Both monkey Z (13 sessions, Wilcoxon rank-sum test, p=2.3576e-05) and monkey H (20 sessions, Wilcoxon rank-sum test, p = 7.6714e-05) had significantly higher distributions of population correlations for correct trials than error trials. Bottom shows the differences in the distributions for each n-back condition (rows; left is small n-backs, right is high n-backs). Both small n-backs (Wilcoxon rank-sum test, p=1.1795e-05) and high n-backs (Wilcoxon rank-sum test, p= 0.0088222) were significantly skewed towards larger values for correct trials compared to error trials.

**Figure S5:**
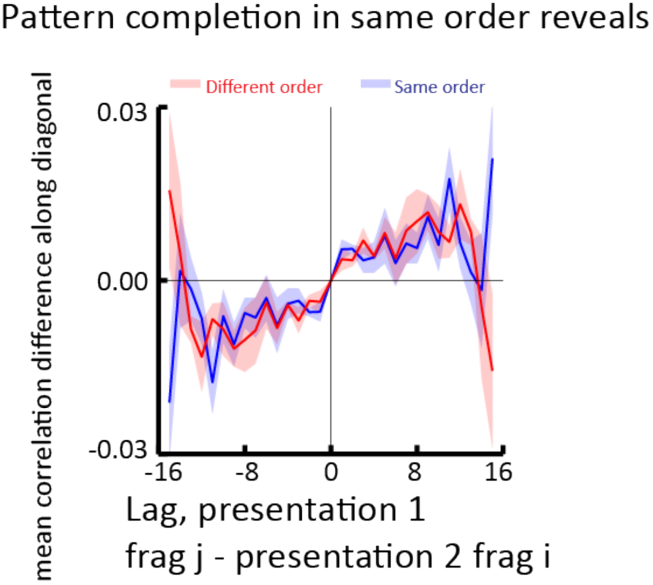
Lag correlations for different image reveal types. Shaded regions indicate SEM. The blue line represents the lag correlation for correct trials in which images were presented in a random but same order on both image presentations. Different images were still presented in different orders. The red line represents the lag correlation for correct trials in which images were presented in a random, different order across both image presentations. Data comes from 5 sessions of monkey Z in which these same and different order reveals were randomly interleaved throughout the session.

**Figure S6:**
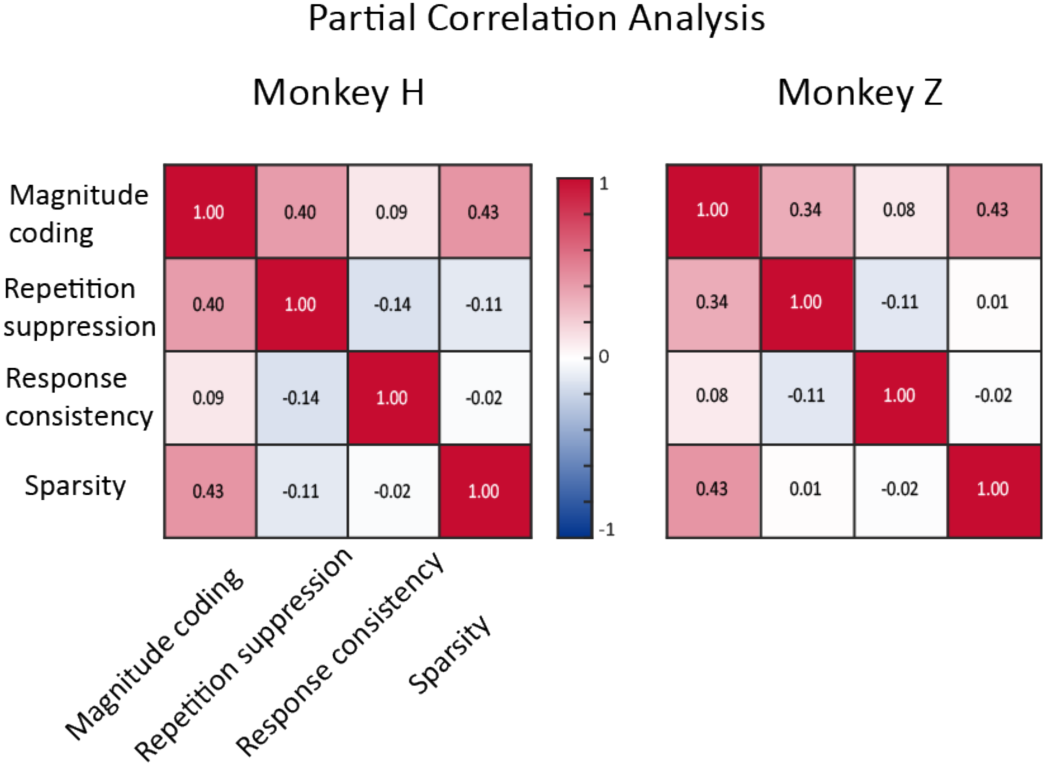
Partial correlation analysis between neural hypotheses. Heatmaps display the linear partial correlation coefficients between the neural components of our hypotheses: repetition suppression (RSI), magnitude coding (mean firing), response consistency (population correlation), and sparsity (proportion of units with zscored firing > 2).

## Bibliography

1. Standing, L. Learning 10000 pictures. Q. J. Exp. Psychol. 25, 207–222 (1973).

2. Meyer, T. & Rust, N. C. Single-exposure visual memory judgments are reflected in inferotemporal cortex. eLife 7, (2018).

3. Shepard, R. N. Recognition memory for words, sentences, and pictures. J. Verbal Learn. Verbal Behav. 6, 156–163 (1967).

4. Vogt, S. & Magnussen, S. Long-term memory for 400 pictures on a common theme. Exp. Psychol. 54, 298–303 (2007).

5. Standing, L., Conezio, J. & Haber, R. N. Perception and memory for pictures: Single-trial learning of 2500 visual stimuli. Psychon. Sci. 19, 73–74 (1970).

6. Brady, T. F., Konkle, T., Alvarez, G. A. & Oliva, A. Visual long-term memory has a massive storage capacity for object details. Proc. Natl. Acad. Sci. 105, 14325–14329 (2008).

7. Cohen, M. R. & Kohn, A. Measuring and interpreting neuronal correlations. Nat. Neurosci. 14, 811–819 (2011).

8. Cohen, M. R. & Maunsell, J. H. R. Attention improves performance primarily by reducing interneuronal correlations. Nat. Neurosci. 12, 1594–1600 (2009).

9. Gregoriou, G. G., Rossi, A. F., Ungerleider, L. G. & Desimone, R. Lesions of prefrontal cortex reduce attentional modulation of neuronal responses and synchrony in V4. Nat. Neurosci. 17, 1003–1011 (2014).

10. Gu, Y. et al. Perceptual Learning Reduces Interneuronal Correlations in Macaque Visual Cortex. Neuron 71, 750–761 (2011).

11. Herrero, J. L., Gieselmann, M. A., Sanayei, M. & Thiele, A. Attention-Induced Variance and Noise Correlation Reduction in Macaque V1 Is Mediated by NMDA Receptors. Neuron 78, 729–739 (2013).

12. Poort, J. et al. Learning and attention increase visual response selectivity through distinct mechanisms. Neuron 110, 686–697.e6 (2022).

13. Luo, T. Z. & Maunsell, J. H. R. Neuronal Modulations in Visual Cortex Are Associated with Only One of Multiple Components of Attention. Neuron 86, 1182–1188 (2015).

14. Mayo, J. P. & Maunsell, J. H. R. Graded Neuronal Modulations Related to Visual Spatial Attention. J. Neurosci. 36, 5353–5361 (2016).

15. Mitchell, J. F., Sundberg, K. A. & Reynolds, J. H. Spatial Attention Decorrelates Intrinsic Activity Fluctuations in Macaque Area V4. Neuron 63, 879–888 (2009).

16. Ni, A. M., Ruff, D. A., Alberts, J. J., Symmonds, J. & Cohen, M. R. Learning and attention reveal a general relationship between population activity and behavior. Science 359, 463– 465 (2018).

17. Mehrpour, V., Meyer, T., Simoncelli, E. P. & Rust, N. C. Pinpointing the neural signatures of single-exposure visual recognition memory. Proc. Natl. Acad. Sci. 118, (2021).

18. Rust, N. C. & Mehrpour, V. Understanding Image Memorability. Trends Cogn. Sci. 24, 557– 568 (2020).

19. Bainbridge, W. A. & Rissman, J. Dissociating neural markers of stimulus memorability and subjective recognition during episodic retrieval. Sci. Rep. 8, (2018).

20. Jaegle, A. et al. Population response magnitude variation in inferotemporal cortex predicts image memorability. eLife 8, e47596 (2019).

21. Mohan, K. et al. Visual familiarity learning at multiple timescales in the primate inferotemporal cortex. Preprint at 10.1101/2024.01.05.574412 (2024).

22. Fahy, F. L., Riches, I. P. & Brown, M. W. Neuronal activity related to visual recognition memory: long-term memory and the encoding of recency and familiarity information in the primate anterior and medial inferior temporal and rhinal cortex. Exp. Brain Res. 96, (1993).

23. Grill-Spector, K. et al. Differential processing of objects under various viewing conditions in the human lateral occipital complex. Neuron 24, 187–203 (1999).

24. Grill-Spector, K., Henson, R. & Martin, A. Repetition and the brain: neural models of stimulus-specific effects. Trends Cogn. Sci. 10, 14–23 (2006).

25. Vidal, J. R. et al. Neural repetition suppression in ventral occipito-temporal cortex occurs during conscious and unconscious processing of frequent stimuli. NeuroImage 95, 129–135 (2014).

26. DiCarlo, J. J. & Cox, D. D. Untangling invariant object recognition. Trends Cogn. Sci. 11, 333–341 (2007).

27. Afraz, A., Yamins, D. L. K. & DiCarlo, J. J. Neural Mechanisms Underlying Visual Object Recognition. Cold Spring Harb. Symp. Quant. Biol. 79, 99–107 (2014).

28. Desimone, R. & Duncan, J. Neural mechanisms of selective visual attention. Annu. Rev. Neurosci. 18, 193–222 (1995).

29. Reynolds, J. H. & Chelazzi, L. Attentional modulation of visual processing. Annu. Rev. Neurosci. 27, 611–647 (2004).

30. Maunsell, J. H. R. Neuronal Mechanisms of Visual Attention. Annu. Rev. Vis. Sci. 1, 373– 391 (2015).

31. Rust, N. C. & Cohen, M. R. Priority coding in the visual system. Nat. Rev. Neurosci. 23, 376–388 (2022).

32. Goetschalckx, L., Andonian, A., Oliva, A. & Isola, P. GANalyze: Toward Visual Definitions of Cognitive Image Properties.

33. Lin, Q., Yousif, S. R., Chun, M. M. & Scholl, B. J. Visual memorability in the absence of semantic content. Cognition 212, 104714 (2021).

34. Pasupathy, A., Popovkina, D. V. & Kim, T. Visual Functions of Primate Area V4. Annu. Rev. Vis. Sci. 6, 363–385 (2020).

35. Roe, A. W. et al. Toward a unified theory of visual area V4. Neuron 74, 12–29 (2012).

36. Miller, E. K. & Cohen, J. D. An integrative theory of prefrontal cortex function. Annu. Rev. Neurosci. 24, 167–202 (2001).

37. Noudoost, B., Chang, M. H., Steinmetz, N. A. & Moore, T. Top-down control of visual attention. Curr. Opin. Neurobiol. 20, 183–190 (2010).

38. Gilbert, C. D. & Li, W. Top-down influences on visual processing. Nat. Rev. Neurosci. 14, 350–363 (2013).

39. Fiebelkorn, I. C. & Kastner, S. Functional Specialization in the Attention Network. Annu. Rev. Psychol. 71, 221–249 (2020).

40. Hayden, B. Y. & Gallant, J. L. Working memory and decision processes in visual area v4. Front. Neurosci. 7, 18 (2013).

41. Motter, B. C. Modulation of Transient and Sustained Response Components of V4 Neurons by Temporal Crowding in Flashed Stimulus Sequences. J. Neurosci. 26, 9683–9694 (2006).

42. Rolls, E. T. The mechanisms for pattern completion and pattern separation in the hippocampus. Front. Syst. Neurosci. 7, 74 (2013).

43. Marr, D. Simple memory: a theory for archicortex. Philos. Trans. R. Soc. Lond. B. Biol. Sci. 262, 23–81 (1971).

44. Mizumori, S. J., McNaughton, B. L., Barnes, C. A. & Fox, K. B. Preserved spatial coding in hippocampal CA1 pyramidal cells during reversible suppression of CA3c output: evidence for pattern completion in hippocampus. J. Neurosci. Off. J. Soc. Neurosci. 9, 3915–3928 (1989).

45. Rolls, E. T. Pattern separation, completion, and categorisation in the hippocampus and neocortex. Neurobiol. Learn. Mem. 129, 4–28 (2016).

46. McNaughton, B. L. & Morris, R. G. M. Hippocampal synaptic enhancement and information storage within a distributed memory system. Trends Neurosci. 10, 408–415 (1987).

47. Neunuebel, J. P. & Knierim, J. J. CA3 Retrieves Coherent Representations from Degraded Input: Direct Evidence for CA3 Pattern Completion and Dentate Gyrus Pattern Separation. Neuron 81, 416–427 (2014).

48. Treves, A. & Rolls, E. T. Computational analysis of the role of the hippocampus in memory. Hippocampus 4, 374–391 (1994).

49. Gold, J. I. & Shadlen, M. N. The Neural Basis of Decision Making. Annu. Rev. Neurosci. 30, 535–574 (2007).

50. Kramer, M. A., Hebart, M. N., Baker, C. I. & Bainbridge, W. A. The features underlying the memorability of objects. Sci. Adv. 9, (2023).

51. Hebart, M. N. et al. THINGS: A database of 1,854 object concepts and more than 26,000 naturalistic object images. PLOS ONE 14, e0223792 (2019).

52. Jutras, M. J. & Buffalo, E. A. Recognition memory signals in the macaque hippocampus. Proc. Natl. Acad. Sci. U. S. A. 107, 401–406 (2010).

53. Jannuzi, B. G. L. et al. Sharpened visual memory representations are reflected in inferotemporal cortex. Preprint at 10.1101/2025.04.28.651105 (2025).

54. Vinje, W. E. & Gallant, J. L. Sparse coding and decorrelation in primary visual cortex during natural vision. Science 287, 1273–1276 (2000).

55. Olshausen, B. A. & Field, D. J. Sparse coding of sensory inputs. Curr. Opin. Neurobiol. 14, 481–487 (2004).

56. Zhou, Z. & Schapiro, A. C. A gradient of complementary learning systems emerges through meta-learning.

57. Leutgeb, J. K. & Moser, E. I. Enigmas of the dentate gyrus. Neuron 55, 176–178 (2007).

58. Xue, G. et al. Greater neural pattern similarity across repetitions is associated with better memory. Science 330, 97–101 (2010).

59. Ward, E. J., Chun, M. M. & Kuhl, B. A. Repetition suppression and multi-voxel pattern similarity differentially track implicit and explicit visual memory. J. Neurosci. Off. J. Soc. Neurosci. 33, 14749–14757 (2013).

60. LaRocque, K. F. et al. Global similarity and pattern separation in the human medial temporal lobe predict subsequent memory. J. Neurosci. Off. J. Soc. Neurosci. 33, 5466–5474 (2013).

61. Hopfield, J. J. Neural networks and physical systems with emergent collective computational abilities. Proc. Natl. Acad. Sci. U. S. A. 79, 2554–2558 (1982).

62. Rolls, E. & Treves, A. Neural Networks and Brain Function. (Oxford University Press, 1997). doi:10.1093/acprof:oso/9780198524328.001.0001.

63. Shepard, R. N. & Teghtsoonian, M. Retention of information under conditions approaching a steady state. J. Exp. Psychol. 62, 302–309 (1961).

64. Hockley, W. E. Retrieval processes in continuous recognition. J. Exp. Psychol. Learn. Mem. Cogn. 8, 497–512 (1982).

65. Mohsenzadeh, Y., Mullin, C., Oliva, A. & Pantazis, D. The perceptual neural trace of memorable unseen scenes. Sci. Rep. 9, (2019).

66. Hebscher, M., Kragel, J. E., Kahnt, T. & Voss, J. L. Enhanced reinstatement of naturalistic event memories due to hippocampal-network-targeted stimulation. Curr. Biol. 31, 1428–1437.e5 (2021).

67. Bainbridge, W. A., Dilks, D. D. & Oliva, A. Memorability: A stimulus-driven perceptual neural signature distinctive from memory. NeuroImage 149, 141–152 (2017).

68. Auksztulewicz, R. & Friston, K. Repetition suppression and its contextual determinants in predictive coding. Cortex J. Devoted Study Nerv. Syst. Behav. 80, 125–140 (2016).

69. Larsson, J. & Smith, A. T. fMRI repetition suppression: neuronal adaptation or stimulus expectation? Cereb. Cortex N. Y. N 1991 22, 567–576 (2012).

70. Williams, N. P. & Olson, C. R. Contribution of individual features to repetition suppression in macaque inferotemporal cortex. J. Neurophysiol. 128, 378–394 (2022).

71. Hsu, Y.-F., Hämäläinen, J. A. & Waszak, F. Repetition suppression comprises both attention-independent and attention-dependent processes. NeuroImage 98, 168–175 (2014).

72. Solomon, S. G. & Kohn, A. Moving sensory adaptation beyond suppressive effects in single neurons. Curr. Biol. CB 24, R1012–1022 (2014).

73. Kohn, A. Visual adaptation: physiology, mechanisms, and functional benefits. J. Neurophysiol. 97, 3155–3164 (2007).

74. Staresina, B. P., Henson, R. N. A., Kriegeskorte, N. & Alink, A. Episodic reinstatement in the medial temporal lobe. J. Neurosci. Off. J. Soc. Neurosci. 32, 18150–18156 (2012).

75. Polyn, S. M., Natu, V. S., Cohen, J. D. & Norman, K. A. Category-specific cortical activity precedes retrieval during memory search. Science 310, 1963–1966 (2005).

76. Johnson, J. D., McDuff, S. G. R., Rugg, M. D. & Norman, K. A. Recollection, familiarity, and cortical reinstatement: a multivoxel pattern analysis. Neuron 63, 697–708 (2009).

77. Gordon, A. M., Rissman, J., Kiani, R. & Wagner, A. D. Cortical reinstatement mediates the relationship between content-specific encoding activity and subsequent recollection decisions. Cereb. Cortex N. Y. N 1991 24, 3350–3364 (2014).

78. McClelland, J. L., McNaughton, B. L. & O’Reilly, R. C. Why there are complementary learning systems in the hippocampus and neocortex: insights from the successes and failures of connectionist models of learning and memory. Psychol. Rev. 102, 419–457 (1995).

79. Ritchey, M., Wing, E. A., LaBar, K. S. & Cabeza, R. Neural similarity between encoding and retrieval is related to memory via hippocampal interactions. Cereb. Cortex N. Y. N 1991 23, 2818–2828 (2013).

80. Peter, A. et al. Stimulus-specific plasticity of macaque V1 spike rates and gamma. Cell Rep. 37, 110086 (2021).

81. Kohn, A. & Movshon, J. A. Neuronal Adaptation to Visual Motion in Area MT of the Macaque. Neuron 39, 681–691 (2003).

82. Barron, H. C., Garvert, M. M. & Behrens, T. E. J. Repetition suppression: a means to index neural representations using BOLD? Philos. Trans. R. Soc. B Biol. Sci. 371, 20150355 (2016).

83. Movshon, J. A. & Lennie, P. Pattern-selective adaptation in visual cortical neurones. Nature 278, 850–852 (1979).

84. Kuravi, P. & Vogels, R. Effect of adapter duration on repetition suppression in inferior temporal cortex. Sci. Rep. 7, 3162 (2017).

85. Miller, E. K., Gochin, P. M. & Gross, C. G. Habituation-like decrease in the responses of neurons in inferior temporal cortex of the macaque. Vis. Neurosci. 7, 357–362 (1991).

86. Ziv, Y. et al. Long-term dynamics of CA1 hippocampal place codes. Nat. Neurosci. 16, 264– 266 (2013).

87. Geva, N., Deitch, D., Rubin, A. & Ziv, Y. Time and experience differentially affect distinct aspects of hippocampal representational drift. Neuron 111, 2357–2366.e5 (2023).

88. Manns, J. R., Howard, M. W. & Eichenbaum, H. Gradual changes in hippocampal activity support remembering the order of events. Neuron 56, 530–540 (2007).

89. Rule, M. E., O’Leary, T. & Harvey, C. D. Causes and consequences of representational drift. Curr. Opin. Neurobiol. 58, 141–147 (2019).

90. Driscoll, L. N., Duncker, L. & Harvey, C. D. Representational drift: Emerging theories for continual learning and experimental future directions. Curr. Opin. Neurobiol. 76, 102609 (2022).

91. Kesner, R. P. & Rolls, E. T. A computational theory of hippocampal function, and tests of the theory: new developments. Neurosci. Biobehav. Rev. 48, 92–147 (2015).

92. Carrillo-Reid, L., Han, S., Yang, W., Akrouh, A. & Yuste, R. Controlling Visually Guided Behavior by Holographic Recalling of Cortical Ensembles. Cell 178, 447–457.e5 (2019).

93. Marshel, J. H. et al. Cortical layer-specific critical dynamics triggering perception. Science 365, eaaw5202 (2019).

94. Bainbridge, W. A., Hall, E. H. & Baker, C. I. Distinct Representational Structure and Localization for Visual Encoding and Recall during Visual Imagery. Cereb. Cortex 31, 1898– 1913 (2021).

95. Tang, H. et al. Spatiotemporal dynamics underlying object completion in human ventral visual cortex. Neuron 83, 736–748 (2014).

96. Rajaei, K., Mohsenzadeh, Y., Ebrahimpour, R. & Khaligh-Razavi, S.-M. Beyond core object recognition: Recurrent processes account for object recognition under occlusion. PLOS Comput. Biol. 15, e1007001 (2019).

97. Tanaka, K. Z. et al. Cortical representations are reinstated by the hippocampus during memory retrieval. Neuron 84, 347–354 (2014).

98. Genkin, M., Shenoy, K. V., Chandrasekaran, C. & Engel, T. A. The dynamics and geometry of choice in the premotor cortex. Nature (2025) doi:10.1038/s41586-025-09199-1.

99. Latimer, K. W., Yates, J. L., Meister, M. L. R., Huk, A. C. & Pillow, J. W. Single-trial spike trains in parietal cortex reveal discrete steps during decision-making. Science 349, 184–187 (2015).

100. Zoltowski, D. M., Latimer, K. W., Yates, J. L., Huk, A. C. & Pillow, J. W. Discrete Stepping and Nonlinear Ramping Dynamics Underlie Spiking Responses of LIP Neurons during Decision-Making. Neuron 102, 1249–1258.e10 (2019).

101. Brunton, B. W., Botvinick, M. M. & Brody, C. D. Rats and Humans Can Optimally Accumulate Evidence for Decision-Making. Science 340, 95–98 (2013).

102. Shadlen, M. N. & Newsome, W. T. Neural basis of a perceptual decision in the parietal cortex (area LIP) of the rhesus monkey. J. Neurophysiol. 86, 1916–1936 (2001).

103. Genkin, M., Hughes, O. & Engel, T. A. Learning non-stationary Langevin dynamics from stochastic observations of latent trajectories. Nat. Commun. 12, 5986 (2021).

104. Srinath, R., Ruff, D. A. & Cohen, M. R. Attention improves information flow between neuronal populations without changing the communication subspace. Curr. Biol. CB 31, 5299–5313.e4 (2021).

105. Ruff, D. A. & Cohen, M. R. Simultaneous multi-area recordings suggest that attention improves performance by reshaping stimulus representations. Nat. Neurosci. 22, 1669– 1676 (2019).

106. Semedo, J. D., Zandvakili, A., Machens, C. K., Yu, B. M. & Kohn, A. Cortical Areas Interact through a Communication Subspace. Neuron 102, 249–259.e4 (2019).

107. Tompary, A., Duncan, K. & Davachi, L. High-resolution investigation of memory-specific reinstatement in the hippocampus and perirhinal cortex. Hippocampus 26, 995–1007 (2016).

108. Gershman, S. J., Fiete, I. & Irie, K. Key-value memory in the brain. Neuron 113, 1694–1707.e1 (2025).

109. Kourtzi, Z. & Kanwisher, N. Representation of perceived object shape by the human lateral occipital complex. Science 293, 1506–1509 (2001).

110. Malach, R. et al. Object-related activity revealed by functional magnetic resonance imaging in human occipital cortex. Proc. Natl. Acad. Sci. U. S. A. 92, 8135–8139 (1995).

111. Orban, G. A., Van Essen, D. & Vanduffel, W. Comparative mapping of higher visual areas in monkeys and humans. Trends Cogn. Sci. 8, 315–324 (2004).

112. Brainard, D. H. The Psychophysics Toolbox. Spat. Vis. 10, 433–436 (1997).

113. Trautmann, E. M. et al. Accurate Estimation of Neural Population Dynamics without Spike Sorting. Neuron 103, 292–308.e4 (2019).

