## SupplementalFigures for "Neuronal signatures of successful one-shot memory in mid-level visual cortex"

**Supplemental**

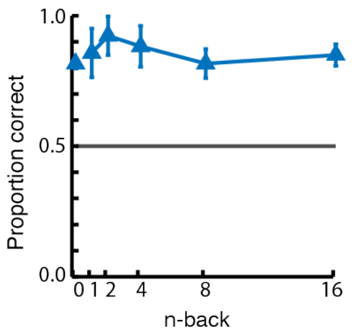


**Figure S1: Behavioral performance on the continuous recognition task without the gradual reveal.** Average accuracy across n-back was computed across three sessions in monkey Z. Error bars show SEM. Performance is better across all n-backs compared to the performance with the gradual reveal in Figure 2c.

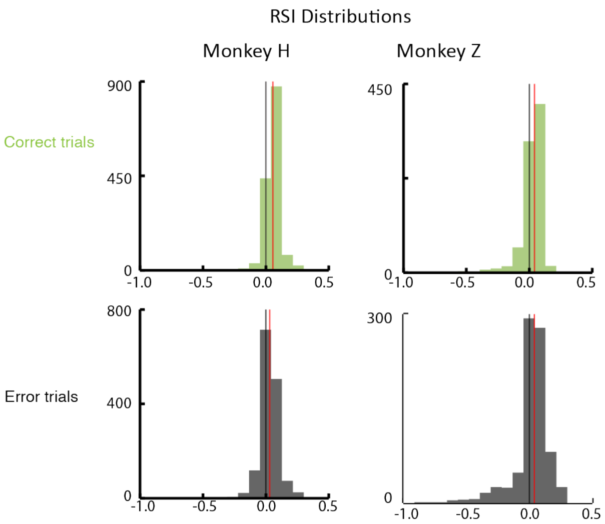


**Figure S2: RSI distributions, split by monkey and behavior.** Columns differentiate monkeys, rows differentiate whether the animal correctly recognized the image or not. Red vertical lines represent median RSI for the respective condition. Black vertical lines mark zero. For every session, we plot the average RSI value derived from each unit (see methods). All distributions were shifted towards positive values, indicating repetition suppression in all monkeys and conditions. Monkey H had significantly more repetition suppression on correct trials compared to errors (rank sum test, 20 sessions,p= 9.1150e-45 ). Monkey Z did not (rank sum test, 13 sessions, p=0.4655).

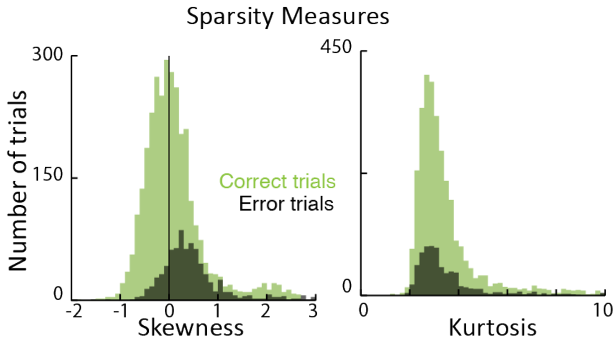


**Figure S3: Sparsity measures.**
Skewness (left) and kurtosis (right) of z-scored firing rate distributions on first image presentations. Color refers to whether the animal was correct (green) or made an error (black) on the next image presentation. No significant difference in the skewness (Wilcoxon rank-sum test, $p = 0.9935$) or kurtosis (Wilcoxon rank-sum test, $p= 0.78957$) for correct versus error trials.

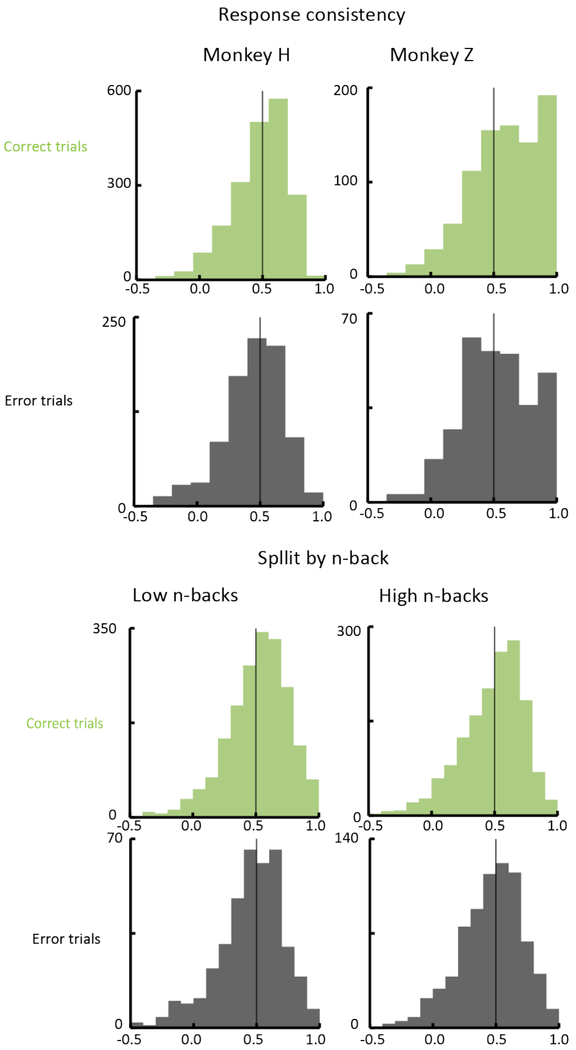


**Figure S4: Population correlation distributions.** Top shows differences in the distributions for each monkey (column) according to whether the animal correctly recognized the image or not on the second presentation (rows; green is correct, black is errors). Both monkey Z (13 sessions, Wilcoxon rank-sum test, p=2.3576e-05) and monkey H (20 sessions, Wilcoxon rank-sum test, p = 7.6714e-05) had significantly higher distributions of population correlations for correct trials than error trials. Bottom shows the differences in the distributions for each n-back condition (rows; left is small n-backs, right is high n-backs). Both small n-backs (Wilcoxon rank-sum test, p=1.1795e-05) and high n-backs (Wilcoxon rank-sum test, p= 0.0088222) were significantly skewed towards larger values for correct trials compared to error trials.

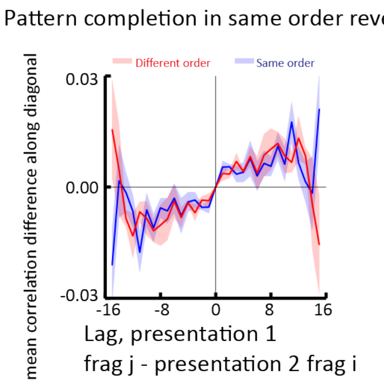


**Figure S5: Lag correlations for different image reveal types.** Shaded regions indicate SEM. The blue line represents the lag correlation for correct trials in which images were presented in a random but same order on both image presentations. Different images were still presented in different orders. The red line represents the lag correlation for correct trials in which images were presented in a random, different order across both image presentations. Data comes from 5 sessions of monkey Z in which these same and different order reveals were randomly interleaved throughout the session.

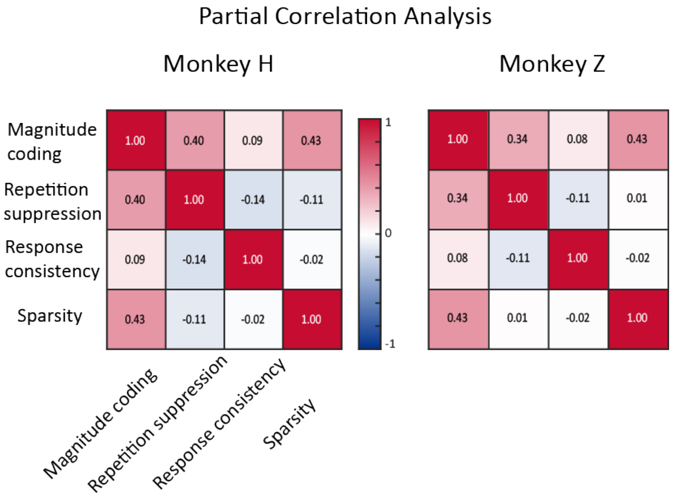


**Figure S6: Partial correlation analysis between neural hypotheses.** Heatmaps display the linear partial correlation coefficients between the neural components of our hypotheses: repetition suppression (RSI), magnitude coding (mean firing), response consistency (population correlation), and sparsity (proportion of units with zscored firing > 2).
